# Hippocampal mechanisms support cortisol-induced memory enhancements

**DOI:** 10.1101/2023.02.08.527745

**Authors:** Brynn E. Sherman, Bailey B. Harris, Nicholas B. Turk-Browne, Rajita Sinha, Elizabeth V. Goldfarb

## Abstract

Stress can powerfully influence episodic memory, often enhancing memory encoding for emotionally salient information. These stress-induced memory enhancements stand at odds with demonstrations that stress and the stress-related hormone cortisol can negatively affect the hippocampus, a brain region important for episodic memory encoding. To resolve this apparent conflict and determine whether and how the hippocampus supports memory encoding under cortisol, we combined behavioral assays of associative memory, high-resolution functional magnetic resonance imaging (fMRI), and pharmacological manipulation of cortisol in a within-participant, double-blinded procedure. Hydrocortisone led to enhanced functional connectivity between hippocampal subregions, which predicted subsequent memory enhancements for emotional information. Cortisol also modified the relationship between hippocampal representations and memory: whereas hippocampal signatures of distinctiveness predicted memory under placebo, relative integration predicted memory under cortisol. Together, these data provide novel evidence that the human hippocampus contains the necessary machinery to support emotional memory enhancements under stress.

## Introduction

Our daily lives are filled with stressful events, from working up against a deadline to hearing that a loved one is ill. Such stressful events can transform the way we encode experiences into memory: Stress can impair memory for neutral, non-stress relevant information (e.g., a conversation with a friend while you were worrying about work), yet enhance memory for emotionally salient or stress-relevant experiences (e.g., a conversation with a family member about their illness) (***Joëls et al., 2006***; ***McGaugh et al., 2015***; ***Shields et al., 2017***; ***Goldfarb, 2019***). Indeed, acute stress can powerfully enhance emotionally salient memories (***Payne et al., 2006, 2007***; ***Smeets et al., 2007***; ***Zoladz et al., 2011***; ***Shields et al., 2022***) and elevated levels of stress-related hormones such as glucocorticoids are associated with enhanced emotional memory in both rodents (***Okuda et al., 2004***; ***Roozendaal et al., 2006***; ***Shors, 2006***; ***Sandi and Pinelo-Nava, 2007***) and humans (***Buchanan and Lovallo, 2001***; ***Abercrombie et al., 2003, 2006***; ***Kuhlmann and Wolf, 2006***; ***Schwabe et al., 2008***; ***Segal et al., 2014***). However, pinpointing the neural mechanisms underlying this selective strengthening of emotional memories under stress presents a puzzle.

Here we focus on the hippocampus, a key region for arbitrating stress effects on memory. The hippocampus plays a critical role in episodic memory by rapidly binding diverse elements of an experience into a detailed, holistic associative memory representation (***Davachi, 2006***). The hippocampus is also highly sensitive to stress, in part because of its high density of glucocorticoid receptors (***Seckl et al., 1991***; ***Lupien et al., 2007***; ***Wang et al., 2013***). Such stress effects are often deleterious: In nonhuman animal models, glucocorticoids tend to impair hippocampal long-term potentiation (LTP), are associated with atrophy of hippocampal neurons, and have been linked to decreased hippocampal neurogenesis (***Kim and Diamond, 2002***). In humans, direct administration of glucocorticoids via hydrocortisone reduces hippocampal BOLD activity (***Pruessner et al., 2008***; ***Lovallo et al., 2010***; ***Bini et al., 2022***). Accordingly, negative stress effects on the hippocampus have been instrumental in explaining stress-induced episodic memory impairments (***McEwen and Sapolsky, 1995***; ***Kim and Diamond, 2002***). Yet, as discussed earlier, stress can also *enhance* memory, including promoting the very types of episodic memory that are thought to be supported by the hippocampus (***van Ast et al., 2014***; ***Goldfarb et al., 2019***). How could stress enhance emotional episodic memories while impairing the function of the brain region important for episodic memory?

One possibility is that stress alters the underlying neural mechanisms of memory encoding, such that memories enhanced under stress rely on extrahippocampal mechanisms. For example, whereas the hippocampus often shows greater BOLD activity for subsequently remembered (versus forgotten) associations (***Davachi et al., 2003***), stress can attenuate (***Qin et al., 2012***) or reverse (***Henckens et al., 2009***) this effect. Stress also alters the electrophysiological signatures of memory encoding (***Meier et al., 2020***). Such findings have been taken as evidence that stress biases memory encoding away from the hippocampus and towards cortical regions, perhaps explaining why memories encoded under stress are less detailed, or more gist-like (***Qin et al., 2012***; ***Pedraza et al., 2016***). Glucocorticoids and stress also broadly influence the neural systems that support memory (***Segal et al., 2010***; ***Hermans et al., 2014***; ***Goldfarb and Phelps, 2017***; ***Schwabe et al., 2022***). Many emotional memory enhancements under stress have been explained by enhanced amygdala activity (***Roozendaal et al., 2009***), as well as hippocampal-amygdala interactions (***Roozendaal and McGaugh, 1997***; ***Kim et al., 2001***; ***Ghosh et al., 2013***; ***Vaisvaser et al., 2013***; ***de Voogd et al., 2017***).

Another possibility is that stress acts directly on hippocampal learning pathways to enhance later memory. Although we mentioned above that glucocorticoids can impair hippocampal LTP, there is also abundant and opposite evidence that they can enhance hippocampal LTP (***Rey et al., 1994***; ***Pavlides et al., 1995***; ***Krugers et al., 2005***; ***Karst and Joëls, 2005***; ***Karst et al., 2005***; ***Vandael et al., 2021***). Such benefits are particularly apparent when stress occurs at the time of synaptic stimulation (i.e., encoding) (***Joëls et al., 2006***; ***Wiegert et al., 2006***). This facilitation is consistent with the role of the hippocampus in regulating stress responses; if stress fully blocked hippocampal function, such regulation would be impossible (***Ulrich-Lai and Herman, 2009***). In humans, cortisol effects on hippocampal BOLD activity have been mixed (***Harrewijn et al., 2020***); in addition to findings of decreased activity, there is also evidence for increased hippocampal BOLD following hydrocortisone (***Symonds et al., 2012***). Additionally, during an acute stress provocation, higher cortisol responses tracked greater hippocampal BOLD increases (***Sinha et al., 2016***). Emotional memory enhancements under stress have also been linked to increased memory-related oscillations in the medial temporal lobe (***Heinbockel et al., 2021***). Together, these data suggest that there may be intra-hippocampal mechanisms to account for emotional memory enhancements with glucocorticoids.

Here we combine high-resolution fMRI, behavioral measures, and double-blind hydrocortisone administration to probe the hippocampal mechanisms underlying memory encoding under cortisol. By using high-resolution fMRI, we embrace the heterogeneity of the hippocampus by examining responses and connectivity profiles of different subfields (e.g., CA1 and a combined CA2/3/dentate gyrus [DG] region). This is a critical advance as nonhuman animal findings demonstrate distinct stress effects across subfields (***Sharvit et al., 2015***; ***Alkadhi, 2019***) and human research indicates that subfields support distinct aspects of memory (***Schapiro et al., 2017***; ***Duncan and Schlichting, 2018***), with CA3 and DG supporting the computations necessary for episodic memory (***Bakker et al., 2008***; ***Molitor et al., 2021***; ***Wanjia et al., 2021***; ***Wammes et al., 2022***). During the fMRI scan, participants encoded emotional and neutral object-scene associations; they then returned to the lab 24h later for memory tests (Figure 1). The emotional objects consisted of alcoholic beverages that generally have positive valence and have been used in prior studies (***Goldfarb et al., 2020***). Participants completed this procedure twice, once receiving 20mg hydrocortisone and once receiving placebo prior to encoding (double-blinded; order counter-balanced). By examining how cortisol influences behavioral measures of memory, neural signatures of encoding within the hippocampus, and brain-behavior relationships, we identify novel mechanisms by which glucocorticoids modulate hippocampal function to enhance emotional memory.

**Figure 1.**
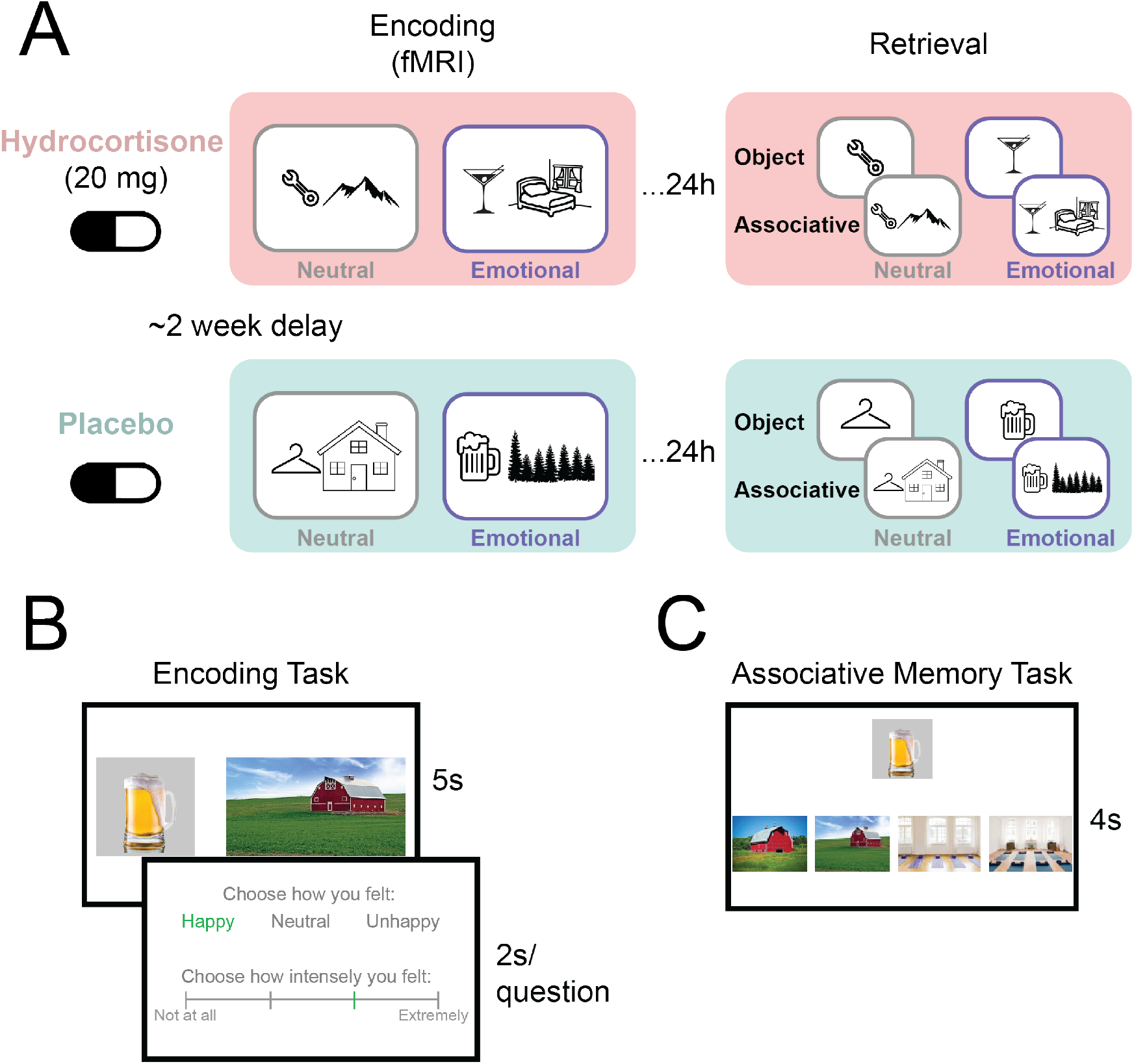
Task design. A) Participants completed the experimental procedure twice (on two separate weeks). They first received a pill containing either 20mg hydrocortisone or no active substance (placebo). They then completed a memory encoding task while undergoing fMRI. The encoding task consisted of two runs of encoding object-scene pair associations. Objects were either neutral (handheld objects) or emotionally salient (alcoholic beverages). Participants returned 24h later to be tested on their memory for individual objects and object-scene associations. B) During encoding, participants viewed an object-scene pair for 5s, during which they imagined the object and scene interacting. They then rated whether the imagined interaction made them feel happy, neutral, or unhappy (valence) and how intensely they felt that way (arousal). C) During the associative memory task, participants were shown an object and asked to identify its associated scene. The options included the correct scene, a perceptually matched lure scene, another scene (which had been encoded with a different object), and a perceptually matched lure for the incorrect scene. Choosing the correct scene denotes correct associative memory.

## Results

### Hydrocortisone administration leads to to elevated cortisol, but no detectable changes in affect or awareness

To validate the hydrocortisone tablet administration, we collected salivary samples throughout the session (Supplemental Figure S1). Indeed, participants exhibited elevated salivary cortisol following drug, but not placebo; drug-induced cortisol elevation was evident throughout encoding (pre- and post-encoding following hydrocortisone relative to placebo: *ps* < 0.001).

Despite this robust increase in peripheral cortisol, we did not observe significant changes in awareness or overall affect (PANAS, measured pre- and post-scan; ***Watson et al., 1988***). Overall, participants were unaware of which pill they had received (immediately post-pill: 9.4% correct, 74% unsure; post-scan: 29% correct, 45% unsure). Furthermore, we found no significant changes in positive [*F* (1, 23) = 0.81, *p* = 0.38] or negative [*F* (1, 23) = 1.39, *p* = 0.25] affect.

### Hydrocortisone modulates subjective affect at encoding

After pill administration, participants completed two runs of an associative memory encoding task while undergoing fMRI (Figure 1A, left). To examine effects of hydrocortisone on emotional memory, each run involved either emotionally salient or neutral trials. One run consisted of forming trial-unique associations between neutral, handheld household objects (e.g., a wrench) and neutral scenes; the other run consisted of associations between an emotionally salient, alcoholic beverages (e.g., a martini) and scenes (more details on stimuli in Methods: Task Stimuli). Participants were instructed to vividly imagine each object and scene interacting and then rate whether the imagined interaction was happy, neutral, or unhappy (valence rating) and how intensely they felt that way (arousal rating; Figure 1B).

Hydrocortisone did not influence arousal ratings, *F* (1, 75) = 0.37, *p* = 0.54. There was a main effect of trial type (emotionally salient vs. neutral) [*F* (1, 75) = 6.27, *p* = 0.015], though this did not interact with pill [*F* (1, 75) = 0.64, *p* = 0.43]. Thus, consistent with the stimulus design, participants rated emotionally salient associations as more arousing than the neutral associations (Figure 2A), and this was not modulated by hydrocortisone.

**Figure 2.**
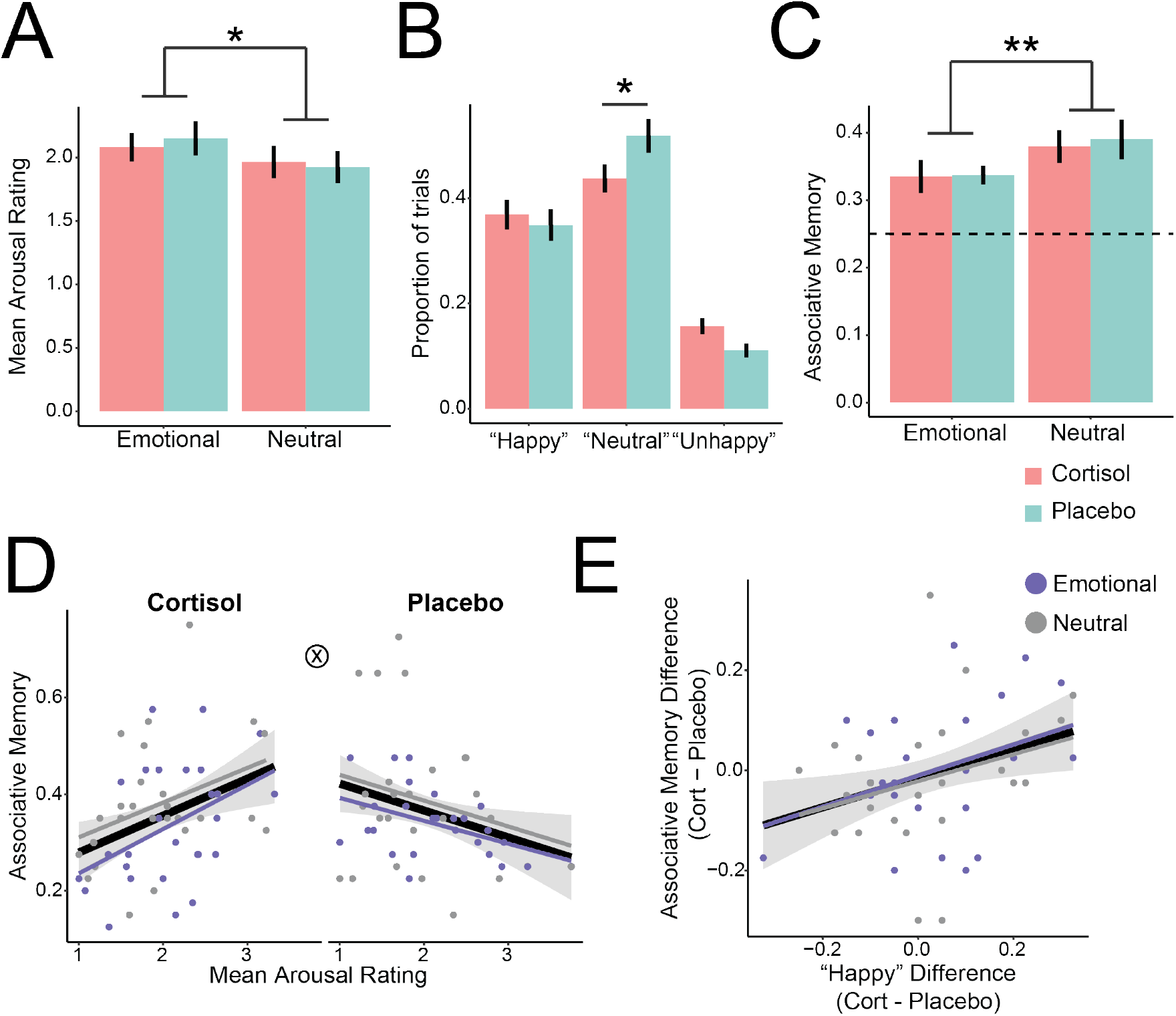
Behavioral Results. A) Participants rated emotionally salient associations as more arousing than neutral associations. B) Cortisol altered the valence of encoded associations, such that participants were less likely to endorse feeling “neutral” about the object-scene pair. C) Participants performed above chance (.25; dashed line) on the associative memory test, with better memory for neutral associations. D) Cortisol altered the relationship between arousal and memory. Participants with high subjective arousal had better associative memories under cortisol, but worse memories under placebo. E) The cortisol-induced change in happiness ratings predicted associative memory, such that participants with greater increases in happiness had better memory. A-C: Error bars denote standard error of the mean across participants; D-E: Error shading indicates 95% confidence interval around the line of best fit, collapsed across trial types (black line). Individual colored lines indicate the line of best fit within each trial type. *p <0.05; **p <0.01

In contrast, hydrocortisone did modulate valence ratings (Figure 2B). There was a main effect of valence on ratings [*F* (2, 279) = 95.11, *p* < 0.001], such that the majority of trials were rated as neutral. There was no main effect of stimulus type [*F* (1, 279) = 0.042, *p* = 0.84], nor did trial type reliably interact with valence [*F* (2, 279) = 2.49, *p* = 0.085]. Although there was no main effect of pill [*F* (1, 279) = 0.062, *p* = 0.80], there was a valence by pill interaction [*F* (2, 279) = 3.53, *p* = 0.031]. This interaction was driven by a smaller proportion of trials rated as “neutral” under hydrocortisone, relative to placebo (*β* = −0.082[*SE* = 0.036], *t*(279) = −2.28, *p* = 0.023). That is, hydrocortisone amplified emotional salience at encoding by shifting participants’ valence ratings away from neutral and towards feeling more positive or negative about the encoded associations.

### Hydrocortisone alters the relationship between arousal and memory encoding

Participants were tested on their memory for encoded associations 24h later. On each trial, participants viewed an object and were asked to select which of four scenes was paired with the object at encoding (Figure 1C). Participants selected the correctly paired scene more often than chance (chance = 0.25; mean proportion correct = 0.36, SD = 0.094). Performance did not differ as a function of pill [*F* (1, 75) = 0.25, *p* = 0.62], though it did differ by trial type [*F* (1, 75) = 8.23, *p* = 0.0054], with better memory for neutral associations (Figure 2C).

Prior work has demonstrated that subjective affect can modulate memory effects under cortisol (***Abercrombie et al., 2003***). Thus, we examined whether memory was affected by subjective arousal differently under hydrocortisone vs. placebo. Indeed, arousal and pill interacted [*F* (1, 71) = 9.30, *p* = 0.0032], such that participants with higher subjective arousal had better associative memory under hydrocortisone, but worse associative memory under placebo (Figure 2D). This difference was significant for both emotional [*β* = 0.080[0.40], *t*(71) = 2.01, *p* = 0.048] and neutral [*β* = 0.095[0.038], *t*(71) = 2.48, *p* = 0.016] trial types.

As hydrocortisone modulated valence ratings during encoding, we asked whether the shifts towards happy or unhappy judgments were related to later memory. Indeed, the change in subjective “happy” ratings predicted the change in associative memory from placebo to hydrocortisone across participants, [*F* (1, 23) = 6.22, *p* = 0.020], with more “happy” ratings under hydrocortisone corresponding to better memory (Figure 2E). This effect did not interact with trial type [*F* (1, 23) = 0.0070, *p* = 0.93] and was specific to positive valence. That is, the change in “unhappy” ratings did not predict differences in memory for either trial type across participants [*F* (1, 23) = 0.36, *p* = 0.56].

Together, these data demonstrate that subjective affect modulates hydrocortisone effects on encoding. Subjective arousal and positive affect bolstered memories encoded with elevated cortisol, but impaired memories under placebo. Although our primary analyses focused on associative memory for object-scene pairs, we observed similar patterns for item recognition of individual objects (Supplemental Figure S2).

### Hydrocortisone enhances intra-hippocampal connectivity to promote emotional memory

After establishing that hydrocortisone interacts with subjective arousal to promote memory, we next sought to investigate whether hippocampal mechanisms could support these memory enhancements. We first examined whether intra-hippocampal connectivity was modulated by hydrocortisone. The hippocampus contains multiple subfields connected to entorhinal cortex (EC), the primary input/output region for the hippocampus (Figure 3A, left). Information from EC is relayed to CA3/DG, which then connects to CA1; CA1 then communicates back out to EC (Figure 3A, right). This circuit (known as the trisynaptic pathway) is particularly important for episodic memory encoding given the sparse connections and high inhibition in CA3 and DG that enable pattern separation (***Schapiro et al., 2017***). We performed a background connectivity analysis (***Al-Aidroos et al., 2012***) to examine how BOLD responses throughout the hippocampal circuit co-fluctuate during encoding following hydrocortisone. Although we ran this analysis for all edges of our simplified hippocampus circuit (EC-CA23DG, CA1-CA23DG, and CA1-EC), we were particularly interested in the CA1-CA23DG edge, as this directly probes hydrocortisone effects on the hippocampus.

**Figure 3.**
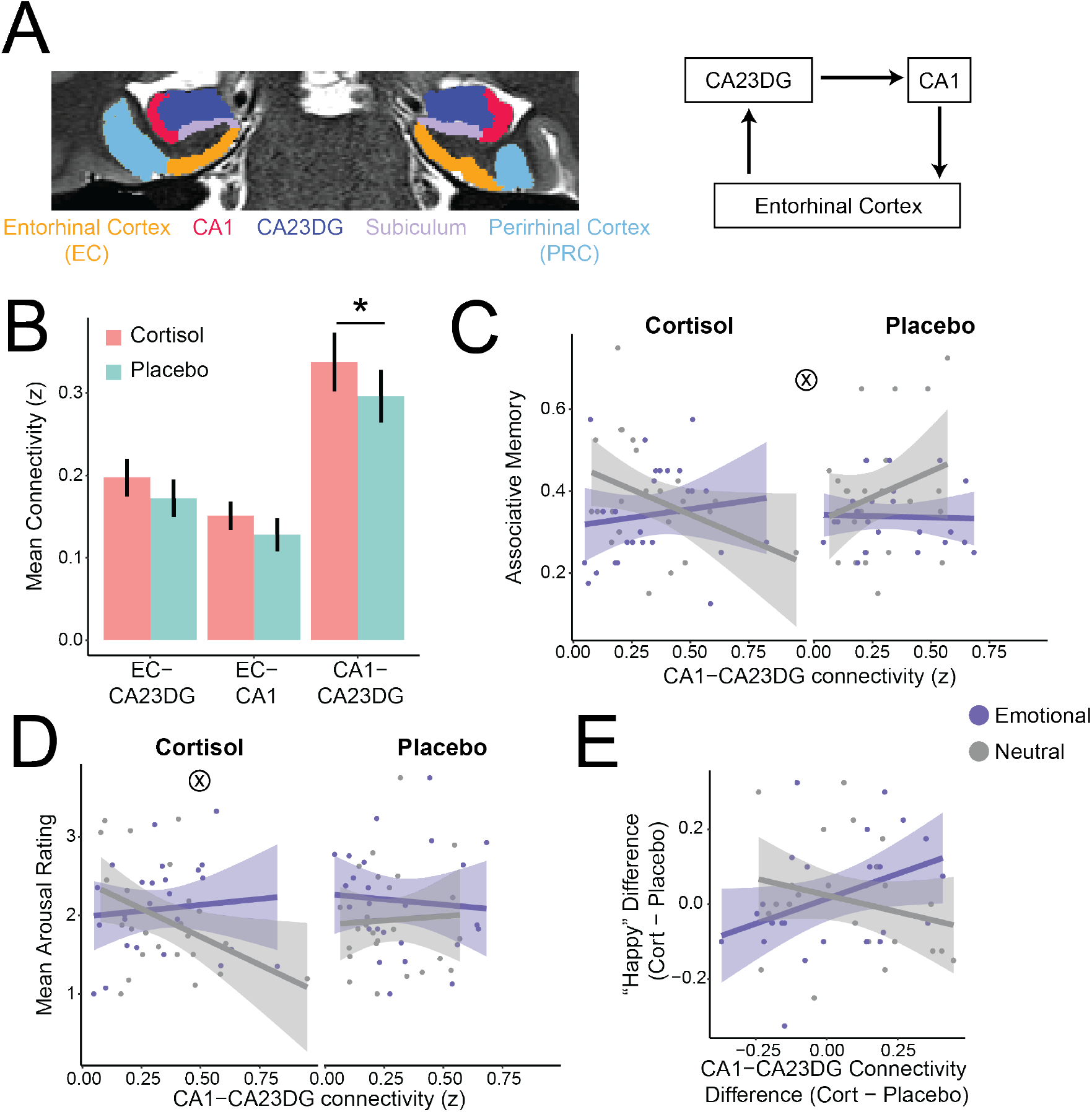
Background Connectivity Results. A) Left: example hippocampal and medial temporal lobe subfields on a representative participant; Right: schematic of hippocampal trisynaptic pathway. B) Hydrocortisone led to increased background connectivity through the hippocampal circuit. C) CA1-CA23DG connectivity interacted with pill and trial type to predict associative memory. D) CA1-CA23DG connectivity interacted with pill to predict subjective arousal. E) The cortisol-induced increase in happiness ratings as a function of the cortisol-induced increase in CA1-CA23DG connectivity. B: Error bars indicates standard error of the mean across participants; *p <0.05. C-E: Error shading indicates 95% confidence interval around the best fit line.

Hydrocortisone enhanced connectivity between CA1 and CA23DG [*F* (1, 72) = 5.20, *p* = 0.026]. Although this effect was only reliable for CA1-CA23DG, it was numerically present for both EC-CA23DG [*F* (1, 72) = 2.01, *p* = 0.16] and EC-CA1 [*F* (1, 72) = 2.38, *p* = 0.13 as well; (Figure 3B)]. Connectivity did not differ between trial types for any of the edges (main effect and cortisol interaction: *ps* > 0.40). These data suggest that cortisol promotes enhanced communication among hippocampal subfields.

Because prior work has suggested that stress can alter hippocampal-amygdala connectivity (e.g., ***Vais-vaser et al., 2013***), we next examined whether cortisol altered connectivity between the amygdala and the hippocampal circuit (Supplemental Figure S3). Hydrocortisone was associated with a marginal decrease in amygdala-EC connectivity [*F* (1, 72) = 3.03, *p* = 0.086], but had no effect on amygdala-CA1 [*F* (1, 72) = 2.21, *p* = 0.14] or amygdala-CA23DG connectivity [*F* (1, 72) = 0.41, *p* = 0.53]. As with intra-hippocampal connectivity, amygdala-hippocampal connectivity did not differ between trial types (main effect and cortisol interaction: *ps* > 0.30). Together, these data suggest a specific role for hydrocortisone in enhancing intra-hippocampal connectivity.

Given that intra-hippocampal, but not amygdala-hippocampal, connectivity was modulated by hydrocortisone, we next tested how intra-hippocampal connectivity related to subsequent associative memory. We observed a three-way interaction between CA1-CA23DG connectivity, trial type, and pill, *F* (1, 68) = 4.92, *p* = 0.030 (Figure 3C). Under placebo, greater intrahippocampal connectivity was associated with stronger memory, and this did not differ between trial types [*β* = −0.16[0.15], *t*(68) = −1.05, *p* = 0.30]. However, following hydrocortisone, greater connectivity positively predicted better memory for emotional, but not neutral associations (emotional vs neutral: *β* = 0.28[0.13], *t*(68) = 2.14, *p* = 0.036).

This finding suggests a reprioritization of hippocampal connectivity to promote emotional, rather than neutral, memories under hydrocortisone. To probe whether this relationship was driven by emotionality, we exploited the positive behavioral relationship between subjective arousal and associative memory with hydrocortisone and assessed whether hippocampal connectivity under hydrocortisone also tracked subjective arousal. Indeed, we observed a significant interaction between connectivity and trial type under hydrocortisone, *F* (1, 23) = 6.54, *p* = 0.018. Mirroring the relationship between connectivity and associative memory, hippocampal connectivity positively tracked arousal for emotional, but not neutral, associations. In contrast, subjective arousal was not related to connectivity under placebo (*ps* > 0.50), although we note that the three-way interaction between connectivity, trial type, and pill was not statistically significant [*F* (1, 68) = 2.03, *p* = 0.16].

We next exploited the behavioral relationship between associative memory and subjective valence, wherein hydrocortisone-induced shifts in positive affect related to better associative memory. We probed the relationship between intra-hippocampal connectivity and subjective valence. We again computed difference scores to assess whether hydrocortisone-associated changes in connectivity related to changes in valence ratings. For positive affect, the difference in CA1-CA23DG connectivity marginally interacted with trial type [*F* (1, 22) = 4.25, *p* = 0.051], such that connectivity changes positively predicted the change in “happy” ratings for emotional, but not neutral trials (Figure 3E). This interaction was not present for negative affect [“unhappy” ratings, *F* (1, 22) = 0.97, *p* = 0.34].

Together, these data suggest an intra-hippocampal mechanism supporting cortisol-induced enhancement of emotional memories. Under placebo, CA1-CA23DG connectivity predicted episodic memory for neutral stimuli, but this relationship shifted with hydrocortisone, with CA1-CA23DG connectivity selectively predicting emotional memory. CA1-CA23DG connectivity also tracked aspects of subjective affect that modulate hydrocortisone effects on associative memory, thereby providing an intra-hippocampal explanation for positive effects of hydrocortisone on emotionally arousing, positively valenced memories.

### Hydrocortisone reverses the relationship between hippocampal pattern similarity and memory

The connectivity approach allowed us to examine how hydrocortisone modulates co-fluctuations in univariate activity between hippocampal subregions. However, the representational content housed in those subregions remains unclear. Prior work using multivariate pattern analysis of hippocampal activity demonstrated that representational distinctiveness, particularly in CA23DG, supports episodic memory encoding (***LaRocque et al., 2013***; ***Wanjia et al., 2021***). This dissimilarity is computationally important, as minimizing overlap between neural patterns associated with similar memories allows those memories to be encoded distinctly, without interference from one another (***Favila et al., 2016***; ***Chanales et al., 2017***). Thus, we next explored the effects of hydrocortisone on representations within these hippocampal subfields during encoding, and how the similarity of these representations relates to memory. We examined within-run pattern similarity, a metric of how similar the hippocampal pattern for an encoded association is to all other associations within that run (Figure 4A).

**Figure 4.**
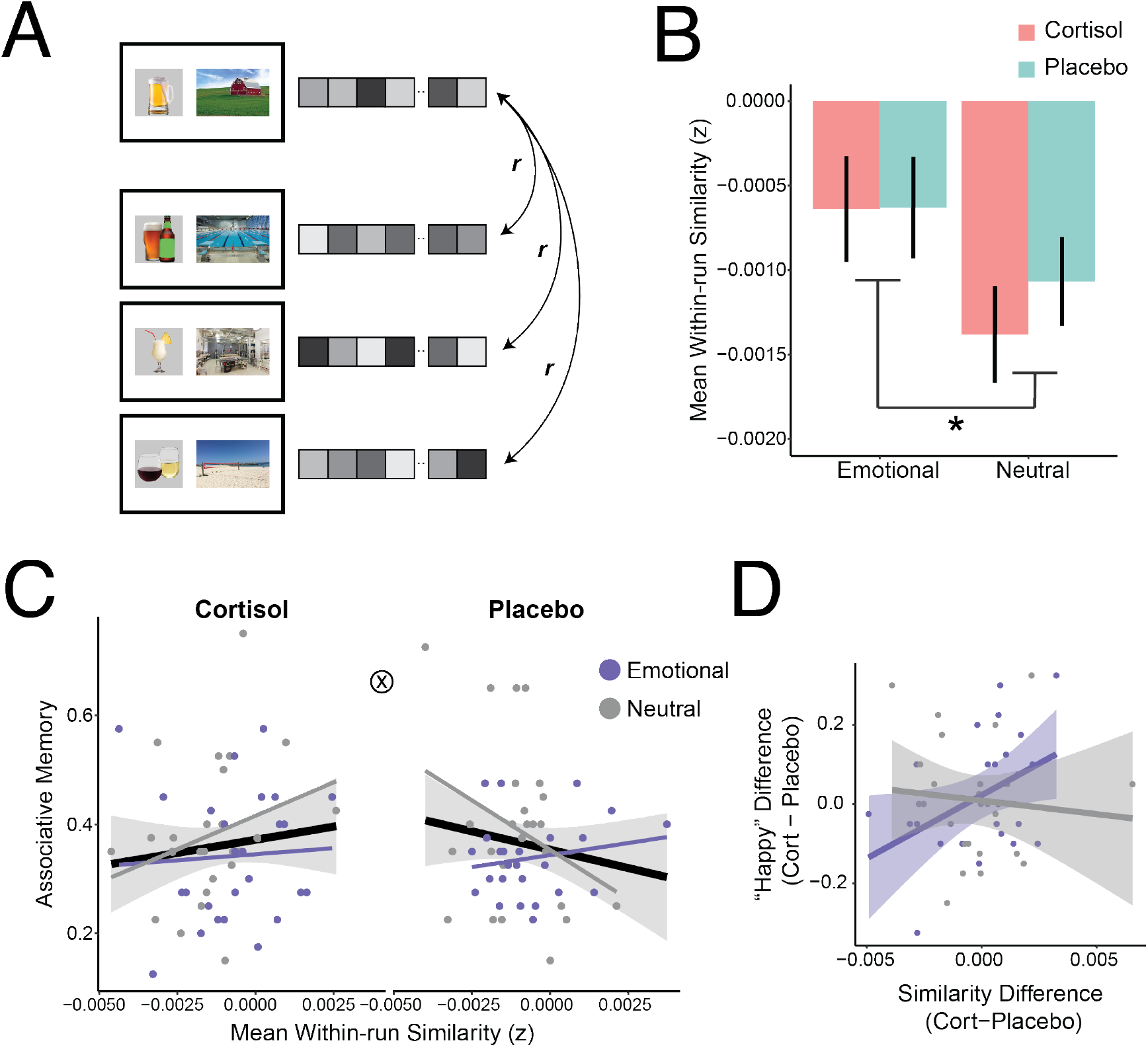
Within-run Pattern Similarity Results. A) We extracted the spatiotemporal pattern associated with each encoding trial. We then correlated each trial to all other trials (averaging the correlations across trials) to obtain a global pattern similarity metric. B) CA23DG pattern similarity differed by trial type, with relatively greater neural similarity among emotional associations. C) CA23DG pattern similarity predicted subsequent associative memory in opposing directions under cortisol versus placebo. D) The difference in CA23DG similarity from cortisol to placebo positively predicted the cortisol-induced increase in happiness ratings from emotional, but not neutral associations. B: Error bars indicate standard error of the mean across participants; *p <0.05. C-D: Error shading indicates 95% confidence interval around the best fit line. Individual colored lines indicate the line of best fit within each trial type.

Hydrocortisone did not affect within-run pattern similarity in either CA1 [*F* (1, 72) = 1.21, *p* = 0.27] or CA23DG [*F* (1, 72) = 0.36, *p* = 0.55]. However, trial type did modulate CA23DG similarity [*F* (1, 72) = 4.86, *p* = 0.031], with relatively greater pattern similarity for emotional, relative to neutral associations (Figure 4B). To ensure that this difference was not due to visual content (i.e., greater inherent visual similarity among emotional objects), we also examined pattern similarity in lateral occipital cortex (LOC), an object-sensitive visual region. There was no effect of either pill [*F* (1, 75) = 0.01, *p* = 0.94] or trial type [*F* (1, 75) = 0.51, *p* = 0.48] on LOC similarity, suggesting that the trial type differences in CA23DG were not driven by visual similarity.

Although there was no pill effect on pattern similarity, the relationship between CA23DG similarity and subsequent associative memory did differ between pills (Figure 4C). Specifically, we observed a similarity by pill interaction [*F* (1, 68) = 5.37, *p* = 0.024], as well as a marginal three-way interaction among similarity, pill, and trial type [*F* (1, 68) = 3.40, *p* = 0.070]. Whereas CA23DG similarity *negatively* predicted memory under placebo (consistent with more distinct neural representations supporting more precise memories), it *positively* predicted memory under hydrocortisone [*β* = 32.2[12.7], *t*(68) = 2.53, *p* = 0.014]. This pattern was not present for CA1, as there was no main effect of pattern similarity on memory, nor interactions with trial type or pill (*ps* > 0.20).

Given that associative memory related to subjective affect, we next examined whether CA23DG similarity predicted arousal or the change in affect ratings. There were no reliable associations between similarity and arousal (*ps* > 0.10). However, CA23DG similarity did relate to the cortisol-induced shift towards positive valence. Increased CA23DG similarity with hydrocortisone tracked increased happiness ratings with hydrocortisone for emotional, but not neutral, associations [*F* (1, 22) = 5.14, *p* = 0.034; Figure 4D].

Together, these data suggest that hydrocortisone reverses the relationship between neural similarity and memory. Consistent with prior work (e.g., ***LaRocque et al., 2013***), CA23DG similarity negatively predicted memory under placebo; this may reflect the computational need for episodic memory to separate memories encoded in a similar temporal context, in order to reduce interference across those memories at test. Intriguingly, however, we observed a positive relationship between similarity and memory under cortisol, suggesting that cortisol may lead memories to be encoded in a fundamentally different, integrated fashion. Relatively greater pattern similarity was also associated with increased “happy” ratings under cortisol, but this effect was specific to emotional associations. Together, these results suggest a distinct hippocampal mechanism for promoting emotionally salient memories under cortisol.

### Hydrocortisone blunts hippocampal subsequent memory effects

Our primary interest in the current study was understanding how intra-hippocampal functional connectivity and representations change with hydrocortisone and contribute to later memory. However, prior work has focused on univariate effects of stress and cortisol, including demonstrating that stress alters hippocampal subsequent memory effects (i.e., greater hippocampal univariate activity for later remembered versus forgotten items; ***Henckens et al., 2009***; ***Qin et al., 2012***). To facilitate comparison with prior work, we thus ran general linear models contrasting subsequently remembered versus forgotten trials (separately for each participant, pill, and trial type; Supplemental Figure S4).

In the whole hippocampus, there was no main effect of pill or trial type, nor an interaction between the two (*ps* > 0.30). However, we observed a numerically positive subsequent memory effects for both trial types under placebo. Collapsing across trial types revealed a reliable subsequent memory effect under placebo [Mean difference = 1.36; SD = 3.24; *t*(24) = 2.10, *p* = 0.046], consistent with prior work (e.g., ***Davachi et al., 2003***). In contrast, although there was not a significant difference between hydrocortisone and placebo [*t*(24) = −0.69, *p* = 0.50], there was no reliable subsequent memory effect for memories encoded under cortisol [Mean difference = 0.38; SD = 4.75, *t*(25) = 0.41, *p* = 0.68]. Considering these effects within hippocampal subfields, we found a similar pattern in CA23DG, but not CA1, consistent prior work and the the proposed role of CA23DG in supporting distinct episodic memories (***Eldridge et al., 2005***; ***Carr et al., 2010***). Together, these results converge with prior demonstrations that stress can alter hippocampal memory formation (***Henckens et al., 2009***; ***Qin et al., 2012***).

## Discussion

In the current study, we combined behavior, high-resolution imaging of the human hippocampus, and pharmacological manipulation of hydrocortisone to provide novel insight into how hippocampal circuitry and representations scaffold memory enhancements under stress. First, we demonstrated behaviorally that cortisol enhances encoding of subjectively emotionally arousing memories. We then demonstrated a role for cortisol in enhancing functional interactions between hippocampal subfields; this intra-hippocampal connectivity supported memory under placebo and selectively supported emotional memory under cortisol. Lastly, we demonstrated that cortisol can alter the way that memories are encoded into the hippocampus, shifting the relationship between neural similiarity and memory from negative under placebo to positive under cortisol. Together, these data provide evidence that mechanisms *within* the hippocampus can support memory enhancements under stress.

Our behavioral findings build on prior demonstrations that subjective affect and cortisol interact to promote memory encoding. Although cortisol administration prior to encoding did not impact memory performance overall, cortisol enhanced memory for participants who experienced greater subjective arousal. This arousal-specific enhancement of memories under stress has been demonstrated previously and highlights the importance of assessing subjective responses to stimuli when measuring stress effects on memory (***Buchanan and Lovallo, 2001***; ***Abercrombie et al., 2003, 2006***; ***Goldfarb et al., 2019***). However, many of these prior studies focused on negative affect (***Abercrombie et al., 2003, 2006***; ***Goldfarb et al., 2019***). By using emotionally salient stimuli (alcoholic beverages) that could be perceived as either positive or negative, we demonstrated that cortisol is particularly beneficial for remembering emotionally arousing, positive episodes. This finding adds to burgeoning literature that acute stress strengthens the formation of positive emotional memories (***Kamp et al., 2019***), and accords with prior work outside the stress domain demonstrating that positive emotion can bolster associative memory (***Madan et al., 2019***). Relatedly, cortisol amplified the perceived emotional salience of encoded memoranda, with participants becoming less likely to rate encoded associations as “neutral” (similar to ***Abercrombie et al., 2003***). Importantly, this cortisol-induced shift in affect valuation was specific to the encoded associations, and did not reflect a broader hydrocortisone-induced change in affect (as shown with PANAS scores). Together, these results are consistent with past studies showing stress and cortisol-induced enhancements in encoding emotionally arousing experiences and provide a key extension by demonstrating the modulatory role of positive affect.

By combining these behavioral metrics with neuroimaging data, we provided insight into hippocampal mechanisms underlying these cortisol-associated memory enhancements. Our use of high-resolution fMRI enabled us to probe the representations of hippocampal subfields, inspired by rodent findings of divergent stress effects across subfields (e.g., ***Alkadhi, 2019***) and human structural imaging results delineating subfield-specific effects of chronic stress and posttraumatic stress disorder (***Wang et al., 2010***; ***Nolan et al., 2020***; ***Weis et al., 2021***). This approach enabled precise localization of hippocampal contributions (rather than aggregating across the whole hippocampus; e.g. ***van Stegeren, 2009***; ***Lovallo et al., 2010***; ***Qin et al., 2012***). Furthermore, using a within-participant pharmacological manipulation, we interrogated the specific role of cortisol in altering encoded memory representations. In doing so, we reveal novel avenues for hydrocortisone to enhance hippocampal function to promote later memory, thus pushing against arguments that stress impairs hippocampal function and shifts memory toward other neural substrates (***Kim and Diamond, 2002***; ***Schwabe et al., 2022***).

We identified a novel role for glucocorticoids in enhancing human hippocampal function: Cortisol was associated with increased communication both among hippocampal subfields. This finding builds on prior rodent work suggesting that stress may alter (***Jacinto et al., 2013***) or enhance (***Stepan et al., 2012***) memory-related theta oscillations within the hippocampus. Importantly, our observed enhancement was specific to the hippocampal circuit; connectivity between the hippocampus and amygdala was not altered by exogenous hydrocortisone. Although this may be surprising given prior work demonstrating that stress enhances hippocampal-amygdala connectivity (***Ghosh et al., 2013***; ***Vaisvaser et al., 2013***), the full stress response includes many processes in addition to cortisol (for example, adrenergic effects on the amygdala play an important role in modulating cortisol effects on the hippocampus; see ***Joëls and Baram, 2009***). Cortisol alone can even reduce hippocampal-amygdala coupling (***Henckens et al., 2012***). This also highlights a limitation in generalizing our focus on cortisol to effects of the multifaceted stress response.

In addition to a broad enhancement with hydrocortisone, we found that intra-hippocampal connectivity related to memory performance under both cortisol and placebo conditions. Under placebo, intra-hippocampal (CA1-CA23DG) connectivity positively predicted memory for neutral associations, consistent with the theorized role for this circuit in supporting episodic memories (***Schapiro et al., 2017***). Under cortisol, in contrast, connectivity positively predicted memory for emotional associations, but negatively for neutral associations. This finding may suggest that the “typical” intrahippocampal mechanism supporting successful memory encoding is repurposed under cortisol to prioritize memory for emotional associations. Highlighting the importance of the emotional nature of these associations, connectivity under cortisol also tracked subjective arousal and valence. Interestingly, the directionality of connectivity/behavior associations differed between emotional and neutral trial types, despite behavioral evidence that arousal broadly tracks memory across trial types. Future work is needed to understand stress and emotion-induced changes in hippocampal encoding mechanisms, and what neural mechanism supports cortisol-induced enhancements in memory for putatively neutral stimuli.

The connectivity results indicate a common mechanism supporting memory: intra-hippocampal connectivity broadly promotes memory under placebo, but selectively promotes emotional memory under cortisol. In contrast, the pattern similarity findings indicate diverging hippocampal encoding processes. Under placebo, pattern *dis*similarity predicted better subsequent memory. This finding is consistent with prior empirical work (***LaRocque et al., 2013***; ***Favila et al., 2016***; ***Chanales et al., 2017***; ***Wanjia et al., 2021***) and theoretical models of hippocampal function, which posit that distinct neural representations are needed to support episodic memory (***McClelland et al., 1995***; ***Brunec et al., 2020***). In contrast, under cortisol, relatively greater pattern *similarity* predicted memory. This pattern is similar to one recently observed in the amygdala (***Bierbrauer et al., 2021***). Closer examination of our data suggest an affect-driven mechanism. First, although the similarity-memory association did not interact with trial type, greater similarity broadly tracked enhanced emotional memory (irrespective of pill). Second, we observed overall greater similarity for emotional compared to neutral stimuli (again, irrespective of pill). This did not appear to be driven by perceptual or semantic features, as this pattern was not observed in lateral occipital cortex. These results may converge with prior demonstrations that greater pattern similarity at encoding (across a range of brain regions, including the hippocampus) predicts better emotional memory (***Visser et al., 2013***; ***Tambini et al., 2017***). Thus, one interpretation of the similarity/memory relationships for neutral memoranda (i.e., negative under placebo but positive under hydrocortisone) is that, with hydrocortisone, emotional encoding mechanisms are engaged to support neutral memory as well. Together, these data suggest that memory formation under cortisol may be supported by distinct underlying computations. Whereas distinctive memory representations may support memory encoding under typical circumstances, more similar memory representations may benefit memory with cortisol.

Considered together, the connectivity and pattern similarity analyses provide evidence that the hippocampus can indeed support enhanced memory formation under hydrocortisone. Importantly, these two signals may serve distinct encoding purposes: intra-hippocampal connectivity primarily explained memory for emotionally salient associations, and CA23DG similarity primarily accounted for memory for neutral information. Importantly, despite robust evidence that the hippocampus can support memory encoding under hydrocortisone, we found evidence for a blunted univariate subsequent memory effect in the hippocampus. Although we interpret these results with caution, given that the difference between placebo and hydrocortisone was not significant, this is consistent with past reports (***Qin et al., 2012***). Although disrupted subsequent memory effects have been interpreted as evidence against hippocampal involvement in stress-induced memory enhancements, our findings challenge this interpretation. Despite replicating this canonical “negative hippocampal” result, we also provide evidence that intrahippocampal function under cortisol can indeed predict subsequent memory. By uncovering positive cortisol effects on hippocampal function, our results highlight the importance of considering multiple hippocampal encoding mechanisms when assessing the effects of cortisol and stress on memory. Whereas cortisol may impair some hippocampal encoding mechanisms, it may yet enhance or alter other avenues by which the hippocampus drives successful memory.

## Acknowledgments

The authors are grateful to Nia Fogelman for assisting with randomization and maintaining the double blind and to Gretchen Hermes for performing physical exams. This work was supported by a Yale Center for Clinical Investigation Junior Faculty Scholar Award (NIH/NCATS KL2 TR001862 to E.V.G.) and the National Institutes of Health (K01 AA027832 to E.V.G.).

## Author contributions

E.V.G. and R.S. conceived the study. E.V.G. and B.B.H. designed the experiments. B.B.H. performed the experiments. E.V.G. and R.S. supervised data collection. N.B.T-B. contributed imaging sequences and analysis pipelines. E.V.G., B.E.S., and N.B.T-B. conceived behavioral and neural analyses. B.B.H. organized and processed the behavioral data. E.V.G. and B.B.H. preprocessed the fMRI data. B.E.S. carried out subsequent analyses on the behavioral and fMRI data. B.E.S. and E.V.G. wrote the manuscript with assistance from B.B.H. All authors edited the manuscript.

## Methods

### Participants

Twenty-seven healthy, right-handed human participants (16 male; mean age 27.6; range 21-44) completed all five sessions of the experiment. This sample size was determined by a power analysis from pilot data indicating that associations between cortisol and memory could be observed with N = 25 (G*Power: correlation = .508, power = 0.85). One participant’s Week 2 data were excluded from all analyses because of technical error (they were shown different stimuli at encoding and retrieval).

Participants were recruited from the New Haven community via online advertisements and flyers. All participants were fluent in English, had BMI between 18-35 (to ensure standardized metabolism of hydrocortisone, which is lipophilic), had normal or corrected-to-normal vision, and did not meet criteria for any substance use disorder (excluding caffeine). To reduce factors that could influence their reaction to hydrocortisone, participants were excluded if they were currently using medications/drugs that interfere with the HPA axis response such as SSRIs, beta-blockers, or corticosteroids. Further, peri- and post-menopausal females, pregnant or lactating females, and those with hysterectomies were excluded. Participants were also excluded based on contraindications for MRI or hydrocortisone tablets. Participants were also required to have had a physical examination within the last 6 months to determine that they could safely complete study procedures; if not, one was administered by a Yale School of Medicine MD. All participants provided written informed consent to complete the study and all procedures were approved by the Yale Institutional Review Board.

### Method Details

#### Procedure Overview

We employed a double-blind, placebo-controlled, crossover design (Figure 1). At the start of each session, participants provided urine samples to undergo drug and pregnancy testing as well as a breathalyzer to ensure sobriety. All experimental sessions occurred between 12:00pm and 6:00pm to control for circadian fluctuations in cortisol (***Lupien et al., 2007***).

After an intake appointment to determine eligibility, participants completed two rounds of encoding (Day 1) and memory retrieval (Day 2, 24h later). These rounds were separated by an average of 16.22 days (SD = 14.53). Prior the encoding session, participants received a tablet containing either 20mg hydrocortisone or a visually identical placebo pill compounded by the Yale Investigational Drug Service (order counterbalanced; see Hydrocortisone Administration and Cortisol Measurement: Randomization). Functional magnetic resonance imaging (fMRI) data were acquired during each encoding session along with salivary samples to measure peripheral cortisol. Participants were instructed not to consume alcohol for 24h prior to fMRI sessions.

#### Intake Appointment

After providing informed consent, participants underwent the Structured Clinical Interview for the DSM-5 or SCID (***First, 2014***) with a trained interviewer to determine if they had ever met criteria for a substance use disorder (SUD) or alcohol use disorder (AUD; N = 3 excluded following intake due to past or present endorsement of AUD). Sobriety was confirmed via breathalyzer at each visit. To further classify drinking behavior, participants completed the AUDIT (***Bush et al., 1998***) administered by an experimenter and answered the Alcohol Severity Index Questionnaire, ASI (***Mäkelä, 2004***), via self-report on an experiment computer.

Participants also self-reported on general demographic information and filled out a series of questionnaires including the Positive Affect Negative Affect Scale, PANAS (***Watson et al., 1988***). If deemed fully eligible at intake, participants were scheduled for the four sessions of the experiment.

#### Tasks

### Encoding

During the fMRI sessions, participants performed an encoding task similar to ***Goldfarb et al. (2020)***. On each trial, participants viewed object and scene photographs and were asked to vividly imagine the object as part of the scene (5s). They then indicated how they felt when imagining each object/scene pair using an MR-compatible button box, reporting their valence (unhappy, happy, or neutral) and arousal (how intensely they felt that way; 1 = not at all; 4 = extremely), and how much they wanted an alcoholic drink (1 = not at all; 4 = a lot; 2s per response). Responses were displayed in green. Trials were separated by a jittered ITI from a geometric distribution (mean = 2s). All task stimuli were presented with MATLAB using the Psychophysics Toolbox (***Brainard, 1997***; ***Pelli, 1997***).

Participants completed two blocks of encoding during each scanning session, and each block contained 40 object-scene pairs. Critically, the two blocks differed in the type of objects presented. One block contained emotionally salient alcohol-related object images (e.g., a glass of wine) and the other block contained neutral, handheld object images (e.g. a tape measure; further details regarding stimuli below). The order of encoding blocks was counterbalanced across participants. Participants were informed that their memory for object-scene pairs would be tested the following day.

### Retrieval

Participants returned 24 hours after each encoding session for a series of memory tests. As stress and cortisol generally impair memory retrieval (***Gagnon and Wagner, 2016***), this timing allowed us to target hydrocortisone effects on memory encoding while avoiding lingering effects on retrieval processes. Memory tests were separated by trial type and occurred in the same order as encoding (i.e., if participants encoded emotional object-scene pairs first, they retrieved emotional memoranda first). No fMRI data were collected at retrieval.

#### Object Recognition

To assess memory for individual objects, participants first viewed all objects from encoding (N = 80) intermixed with novel foils from the same object subcategories (N = 80). After viewing each object (3s), participants indicated whether they thought the image was old (from the encoding session the day before) or new. Responses were on a 4-point scale (“confident old”, “unsure old”, “unsure new”, “confident new”; 2s per response, 0.5s ITI).

#### Associative Memory

To assess memory for the object-scene pairings, participants were shown an object image from encoding. They were first asked if it was paired with an indoor or outdoor scene (maximum response time 2s). They were then shown the same object image along with four scenes (2s): the original scene paired at encoding, a different scene presented at encoding to control for familiarity, and two matched perceptual lures (one lure per encoded scene). Participants indicated with which of the four scenes the object was paired (up to 4s). They were told that if they remembered what scene was shown with the object, but not exactly which image was displayed, to make their best guess between the two images depicting that scene. Pairing the scenes with perceptually matched foils allowed us to dissociate more general, or gist-based memories (e.g., the picture I saw was a beach) versus specific associative memories (e.g., the picture I saw was *that* beach). They were last asked how confident they were in their memory for the scene (1 = “not at all”; 4 = “extremely”; up to 2s). Questions were separated by an ISI of .5s. Choosing the correct, specific scene from encoding denoted correct associative memory.

#### Task Stimuli

##### Objects

A total of 400 photographs of emotionally salient alcohol and neutral handheld stimuli were obtained from prior studies (***Dunsmoor et al., 2012***; ***Fey et al., 2017***; ***Van Der Linden et al., 2015***; ***Sinha et al., 2022***) and from Google image searches. All images were edited to appear on a grey background with visible text occluded and were resized to 200 × 200 pixels. Images were chosen to be perceptually distinct from one another and were evenly distributed into four subcategories (alcohol: beer, wine, liquor, and mixed drinks; neutral: items likely to be found in a kitchen, garage, bathroom, and office). A separate validation experiment was conducted to select a subset of 320 images matched on perceptual (e.g., level of detail (***Dager et al., 2014***) and familiarity (***Bainbridge et al., 2017***)), but not affective features. We chose not to match stimuli on valence or arousal, as we aimed for emotional, but not neutral, stimuli to induce affect.

##### Scenes

A total of 320 indoor and outdoor scene images were obtained from the SUN database (***Xiao et al., 2010***) and Google Image searches. Specifically, we obtained 80 indoor and 80 outdoor scene images, each with a perceptual match (e.g., 2 pictures of beaches) that served as a foil during the associative memory test. As with object stimuli, a separate validation sample was collected to confirm that perceptual similarity across pairmate images was matched.

#### Hydrocortisone Administration and Cortisol Measurement

##### Randomization

After intake, participants were pseudorandomly assigned to receive either hydrocortisone or placebo prior to their first encoding session, taking into account their age, sex, level of education, and drinking level. Pill order was determined by an unblinded statistician, leaving the experimenter (B.B.H.) blind to participant condition.

##### Pill Administration

Across the two weeks, participants received one oral tablet of hydrocortisone 20mg and one oral tablet of placebo (sucrose). The two pills were physically identical. The order of pill administration (week 1 or 2) was counterbalanced by an unblinded statistician, with all additional experimental personnel and participants blinded for the duration of the study. Pills were compounded by the Yale Investigational Drug Service and stored at 20-25C. Pills were administered approximately one hour prior to the start of the first encoding run (consistent with ***Buchanan and Lovallo, 2001***).

##### Measuring cortisol levels

Participants provided six salivary samples over the two fMRI sessions (3 per session) to measure cortisol concentration. The baseline sample was obtained approximately 10 minutes after arrival (after acclimation to the environment and prior to pill administration). The encoding sample was obtained immediately prior to the first encoding run, approximately 1 hour after pill administration. The final sample was obtained after participants exited the scanner. Samples were collected using Starstedt Salivate Tubes and samples were processed by the Yale Center for Clinical Investigation (YCCI) using radioimmunoassay (RIA).

##### Measuring Awareness

To measure subjective awareness of pill administration, participants reported which pill they thought they had received (response options: Cortisol, Placebo, or Not Sure). This was only asked on pill administration (scan) days, both immediately after consuming the pill and after the fMRI scan.

#### fMRI Procedure

Participants underwent fMRI scanning after pill administration on both days. Specifically, participants performed the encoding task (described above) while BOLD fMRI data were acquired. We additionally collected a localizer run and resting-state fMRI scans.

##### Localizer Run

Prior to encoding, participants completed a 6-min run in which they viewed images (1s each, 0.5s ITI) and were instructed to button press anytime an image repeated twice in a row (1-back). They viewed 8 blocks of 22 images each. The blocks consisted of scenes, emotionally salient alcoholic beverages, neutral handheld objects, or phase-scrambled versions of the alcohol and neutral images. None of these images were repeated in the subsequent encoding task. Each category appeared twice during the 8 blocks in a randomized order per subject.

##### Rest Runs

Participants underwent three 6-minute rest scans throughout each session: one prior to encoding (after the localizer run) and one immediately after each encoding run. No data from these rest runs are reported in the current paper.

#### MRI Acquisition Parameters

Data were acquired on Siemens 3T Prisma scanners using a 64-channel coil at the Magnetic Resonance Research Center at Yale University. Data were acquired across three scanners (N = 6 on scanner A, N = 2 on scanner B, N = 19 on scanner C). Each participant completed both of their MRI sessions on a single scanner. Parameters were the same across scanners. Functional images were acquired using an echoplanar imaging (EPI) sequence with the following parameters: TR = 1000ms, TE = 30ms, 75 axial slices, voxel size = 2×2×2 mm, flip angle = 55 degrees, multiband factor = 5, interleaved acquisition, FOV: 220×220.

Anatomical data were acquired using one T1-weighted 3D MPRAGE sequence (TR = 2400ms, TE = 1.22ms, 208 sagittal slices, voxel size = 1×1×1 mm, flip angle = 8 degrees, FOV: 256×256) and one T2-weighted turbo spin echo (TR=11170 ms, TE = 93ms, 54 coronal slices, voxel size = 0.44 × 0.44 × 1.5mm, distance factor=20%, flip angle = 150 degrees).

### Quantification and Statistical Analysis

#### fMRI Preprocessing

fMRI data were preprocessed using FSL 6.0.1. All encoding runs met criteria for inclusion based on motion (defined a priori as >1.5mm absolute mean frame-to-frame displacement, as computed by FSL’s MCLFIRT; ***Jenkinson et al., 2002***). Data were skull-stripped (BET; ***Smith, 2002***), pre-whitened (FILM; ***Woolrich et al., 2001***), and high-pass filtered at 0.01 Hz to remove low-frequency signal drift. We then used FSL’s FEAT () to run a GLM per run to control for motion and covariates of no interest. Regressors included 6 linear estimated motion parameters and white matter timeseries (each plus temporal derivatives) and stick function regressors for nonlinear motion outliers. No smoothing was applied.

For background connectivity analyses (see below), we additionally removed trial-evoked signal (image on/offset and button presses modeled using boxcars convolved with a double-gamma HRF, plus temporal derivatives).

In all analyses, model residuals were aligned to a reference functional scan and then to the participant’s high-resolution T1 anatomical scan using boundary based registration (***Greve and Fischl, 2009***). The highresolution T2 anatomical image (used for defining hippocampal subregions; see below) was also registered to the participant’s T1 anatomical scan using FSL’s FLIRT (***Jenkinson and Smith, 2001***).

#### Regions of Interest Definition

Hippocampal subfields and medial temporal lobe cortical regions were defined individually for each participant primarily based on their T2-weighted anatomical images. Segmentation was done automatically (using both the T1- and T2-weighted anatomical images) using the automated segmentation of hippocampus subfields (ASHS) software package (***Yushkevich et al., 2015***). We used an atlas containing 51 manual segmentations of hippocampal subfields (***Aly and Turk-Browne, 2016a***,b). The automated segmentations were visually inspected for quality assurance and in 4 cases when automatic segmentation was particularly poor, manual segmentation was performed. Manual segmentation was performed using the procedure (i.e., using the same anatomical landmarks) as the segmentations which comprised the atlas (***Insausti et al., 1998***; ***Pruessner et al., 2002***; ***Duvernoy, 2005***), as described in detail in ***Aly and Turk-Browne (2016b)***. The hippocampus was segmented into CA1, CA2/3, and dentate gyrus (DG) subfields; medial temporal lobe cortex was segmented into entrohinal cortex (EC), perirhinal cortex (PRC), and parahippocampal cortex (PHC). For analysis purposes, the CA2/3 and DG subfields were concatenated into a single CA23DG subfield. Further, a whole hippocampus ROI was constructed by concatenating the CA23DG, CA1, and subiculum ROIs. One participant did not have a high-resolution T2-weighted image, and thus their hippocampal subfields could not be segmented; this participant was excluded from all neural analysis looking at the hippocampus.

In addition to MTL ROIs, we analyzed data from amygdala and lateral occipital cortex (LOC). Anatomical amygdala ROIs were defined for each participant based on their T1 MPRAGE scans using FSL’s FIRST automated segmentation tool (***Patenaude et al., 2011***). LOC ROIs were functionally defined based on the localizer scan. These data preprocesed as described above for encoding runs, then residuals were smoothed (6mm FWHM), and entered into subject-level GLMs to extract beta values per block type (emotionally salient objects, neutral objects, scenes, and phase-scrambled images). These subject-level estimates were aligned to MNI space and entered into a group-level ANOVA (AFNI’s 3dANOVA3) as a function of pill and image type. LOC was defined from a post-hoc contrast of neutral objects vs. scrambled images, cluster-corrected (*p* < .001, *α* = .05, AFNI’s 3dClustSim) and then masked with the Harvard-Oxford Probabilistic Atlas definition for LOC (50% threshold). This mask was then aligned to each participant’s functional data.

#### Background Connectivity Analysis

To examine fluctuations among hippocampal subfields during encoding, we conducted a background connectivity analysis (e.g., ***Norman-Haignere et al., 2012***; ***Al-Aidroos et al., 2012***; ***Córdova et al., 2016***). After regressing out the task-evoked signal from each fMRI run as described above, we extracted the mean residual timeseries across voxels in each ROI. We then correlated the timeseries between pairs of ROIs. These correlations were then normalized using a Fisher *r*-to-*z* transform for further analysis.

#### Representational Similarity Analysis

To probe the representational content of encoded associations in the hippocampus, we computed within-run global pattern similarity (similar to ***LaRocque et al., 2013***; ***Tompary and Davachi, 2017***; ***Cowan et al., 2020***). For each encoded association, we extracted the associated pattern of activity per ROI across voxels and time (5 TRs during which the object-scene pair was on screen). To account for the hemodynamic lag, we shifted the data by 5s (5 TRs), such that the extracted pattern reflected the BOLD activity 5-10s after the true onset. We then correlated these spatiotemporal vectors among all pairs of trials within a run and computed the average correlation. As in the background connectivity analyses, we then normalized these averaged correlations via Fisher *r*-to-*z* transform.

#### Univariate Subsequent Memory Analysis

To examine whether hippocampal activation differentiated subsequently remembered versus forgotten associations, we conducted a univariate subsequent memory analysis (e.g., ***Davachi et al., 2003***). We first smoothed the residuals from the preprocessing models above (6mm FWHM) and then ran a separate GLM for each encoding run for each participant. We included separate regressors for subsequently remembered versus forgotten trials plus their temporal derivative. Each trial was modeled as a boxcar (with a duration of 5s), convolved with a double-gamma HRF. We then computed the contrast between subsequently remembered and subsequently forgotten associations. For each ROI, we extracted these contrast estimates averaged across voxels within the ROI, separately for each trial type and pill. We refer to this difference as the “Subsequent Memory Effect”.

#### Statistical Modeling

All statistical analyses were conducted as linear mixed effect models and were performed in R (version 4.1.3) using the nlme package (***Pinheiro et al., 2022***). All models treated participant as a random effect, such that a random intercept was computed for each participant. For analyses that examined the effects of trial type and pill, we additionally included covariates of week, pill order, and trial type order. For analyses that examined a difference between cortisol and placebo, we included covariates of pill order and trial type order. Follow-up tests were performed using the emmeans package (***Lenth, 2022***), with the exception of the subsequent memory effect analyses, in which we used one-sample t-tests to quantify whether remembered versus forgotten contrasts differed from 0.

## References

Abercrombie, H. C., Kalin, N. H., Thurow, M. E., Rosenkranz, M. A., and Davidson, R. J. (2003). Cortisol variation in humans affects memory for emotionally laden and neutral information. Behavioral neuroscience, 117(3):505.

Abercrombie, H. C., Speck, N. S., and Monticelli, R. M. (2006). Endogenous cortisol elevations are related to memory facilitation only in individuals who are emotionally aroused. Psychoneuroendocrinology, 31(2):187–196.

Al-Aidroos, N., Said, C. P., and Turk-Browne, N. B. (2012). Top-down attention switches coupling between low-level and high-level areas of human visual cortex. Proceedings of the National Academy of Sciences, 109(36):14675–14680.

Alkadhi, K. A. (2019). Cellular and molecular differences between area ca1 and the dentate gyrus of the hippocampus. Molecular neurobiology, 56(9):6566–6580.

Aly, M. and Turk-Browne, N. B. (2016a). Attention promotes episodic encoding by stabilizing hippocampal representations. Proceedings of the National Academy of Sciences, 113(4):E420–E429.

Aly, M. and Turk-Browne, N. B. (2016b). Attention stabilizes representations in the human hippocampus. Cerebral Cortex, 26(2):783–796.

Bainbridge, W. A., Dilks, D. D., and Oliva, A. (2017). Memorability: A stimulus-driven perceptual neural signature distinctive from memory. NeuroImage, 149:141–152.

Bakker, A., Kirwan, C. B., Miller, M., and Stark, C. E. (2008). Pattern separation in the human hippocampal ca3 and dentate gyrus. science, 319(5870):1640–1642.

Bierbrauer, A., Fellner, M.-C., Heinen, R., Wolf, O. T., and Axmacher, N. (2021). The memory trace of a stressful episode. Current Biology, 31(23):5204–5213.

Bini, J., Parikh, L., Lacadie, C., Hwang, J. J., Shah, S., Rosenberg, S. B., Seo, D., Lam, K., Hamza, M., De Aguiar, R. B., et al. (2022). Stress-level glucocorticoids increase fasting hunger and decrease cerebral blood flow in regions regulating eating. NeuroImage: Clinical, 36:103202.

Brainard, D. H. (1997). The psychophysics toolbox. Spatial Vision, 10(4):433–436.

Brunec, I. K., Robin, J., Olsen, R. K., Moscovitch, M., and Barense, M. D. (2020). Integration and differentiation of hippocampal memory traces. Neuroscience & Biobehavioral Reviews, 118:196–208.

Buchanan, T. W. and Lovallo, W. R. (2001). Enhanced memory for emotional material following stress-level cortisol treatment in humans. Psychoneuroendocrinology, 26(3):307–317.

Bush, K., Kivlahan, D. R., McDonell, M. B., Fihn, S. D., Bradley, K. A., (ACQUIP, A. C. Q. I. P., et al. (1998). The audit alcohol consumption questions (audit-c): an effective brief screening test for problem drinking. Archives of Internal Medicine, 158(16):1789–1795.

Carr, V. A., Viskontas, I. V., Engel, S. A., and Knowlton, B. J. (2010). Neural activity in the hippocampus and perirhinal cortex during encoding is associated with the durability of episodic memory. Journal of Cognitive Neuroscience, 22(11):2652–2662.

Chanales, A. J., Oza, A., Favila, S. E., and Kuhl, B. A. (2017). Overlap among spatial memories triggers repulsion of hippocampal representations. Current Biology, 27(15):2307–2317.

Córdova, N. I., Tompary, A., and Turk-Browne, N. B. (2016). Attentional modulation of background connectivity between ventral visual cortex and the medial temporal lobe. Neurobiology of learning and memory, 134:115–122.

Cowan, E., Liu, A., Henin, S., Kothare, S., Devinsky, O., and Davachi, L. (2020). Sleep spindles promote the restructuring of memory representations in ventromedial prefrontal cortex through enhanced hippocampal–cortical functional connectivity. Journal of Neuroscience, 40(9):1909–1919.

Dager, A. D., Anderson, B. M., Rosen, R., Khadka, S., Sawyer, B., Jiantonio-Kelly, R. E., Austad, C. S., Raskin, S. A., Tennen, H., Wood, R. M., et al. (2014). Functional magnetic resonance imaging (fmri) response to alcohol pictures predicts subsequent transition to heavy drinking in college students. Addiction, 109(4):585–595.

Davachi, L. (2006). Item, context and relational episodic encoding in humans. Current opinion in neurobiology, 16(6):693–700.

Davachi, L., Mitchell, J. P., and Wagner, A. D. (2003). Multiple routes to memory: distinct medial temporal lobe processes build item and source memories. Proceedings of the National Academy of Sciences, 100(4):2157–2162.

de Voogd, L. D., Klumpers, F., Fernández, G., and Hermans, E. J. (2017). Intrinsic functional connectivity between amygdala and hippocampus during rest predicts enhanced memory under stress. Psychoneuroendocrinology, 75:192–202.

Duncan, K. D. and Schlichting, M. L. (2018). Hippocampal representations as a function of time, subregion, and brain state. Neurobiology of learning and memory, 153:40–56.

Dunsmoor, J. E., Martin, A., and LaBar, K. S. (2012). Role of conceptual knowledge in learning and retention of conditioned fear. Biological psychology, 89(2):300–305.

Duvernoy, H. M. (2005). The Human Hippocampus: Functional Anatomy, Vascular-ization and Serial Sections with MRI. Springer.

Eldridge, L. L., Engel, S. A., Zeineh, M. M., Bookheimer, S. Y., and Knowlton, B. J. (2005). A dissociation of encoding and retrieval processes in the human hippocampus. Journal of Neuroscience, 25(13):3280–3286.

Favila, S. E., Chanales, A. J., and Kuhl, B. A. (2016). Experience-dependent hippocampal pattern differentiation prevents interference during subsequent learning. Nature communications, 7(1):1–10.

Fey, W., Moggi, F., Rohde, K. B., Michel, C., Seitz, A., and Stein, M. (2017). Development of stimulus material for research in alcohol use disorders. International Journal of Methods in Psychiatric Research, 26(1):e1527.

First, M. B. (2014). Structured clinical interview for the dsm (scid). The encyclopedia of clinical psychology, pages 1–6.

Gagnon, S. A. and Wagner, A. D. (2016). Acute stress and episodic memory retrieval: neurobiological mechanisms and behavioral consequences. Annals of the New York Academy of Sciences, 1369(1):55–75.

Ghosh, S., Laxmi, T. R., and Chattarji, S. (2013). Functional connectivity from the amygdala to the hippocampus grows stronger after stress. Journal of Neuroscience, 33(17):7234–7244.

Goldfarb, E. V. (2019). Enhancing memory with stress: progress, challenges, and opportunities. Brain and Cognition, 133:94–105.

Goldfarb, E. V., Fogelman, N., and Sinha, R. (2020). Memory biases in alcohol use disorder: enhanced memory for contexts associated with alcohol prospectively predicts alcohol use outcomes. Neuropsychopharmacology, 45(8):1297–1305.

Goldfarb, E. V. and Phelps, E. A. (2017). Stress and the trade-off between hippocampal and striatal memory. Current Opinion in Behavioral Sciences, 14:47–53.

Goldfarb, E. V., Tompary, A., Davachi, L., and Phelps, E. A. (2019). Acute stress throughout the memory cycle: Diverging effects on associative and item memory. Journal of Experimental Psychology: General, 148(1):13.

Greve, D. N. and Fischl, B. (2009). Accurate and robust brain image alignment using boundary-based registration. Neuroimage, 48(1):63–72.

Harrewijn, A., Vidal-Ribas, P., Clore-Gronenborn, K., Jackson, S. M., Pisano, S., Pine, D. S., and Stringaris, A. (2020). Associations between brain activity and endogenous and exogenous cortisol–a systematic review. Psychoneuroendocrinology, 120:104775.

Heinbockel, H., Quaedflieg, C. W., Schneider, T. R., Engel, A. K., and Schwabe, L. (2021). Stress enhances emotional memory-related theta oscillations in the medial temporal lobe. Neurobiology of Stress, 15:100383.

Henckens, M. J., Hermans, E. J., Pu, Z., Joëls, M., and Fernández, G. (2009). Stressed memories: how acute stress affects memory formation in humans. Journal of Neuroscience, 29(32):10111–10119.

Henckens, M. J., van Wingen, G. A., Joels, M., and Fernandez, G. (2012). Corticosteroid induced decoupling of the amygdala in men. Cerebral cortex, 22(10):2336–2345.

Hermans, E. J., Henckens, M. J., Joëls, M., and Fernández, G. (2014). Dynamic adaptation of large-scale brain networks in response to acute stressors. Trends in neurosciences, 37(6):304–314.

Insausti, R., Juottonen, K., Soininen, H., Insausti, A. M., Partanen, K., Vainio, P., Laakso, M. P., and Pitkänen, A. (1998). Mr volumetric analysis of the human entorhinal, perirhinal, and temporopolar cortices. American journal of neuroradiology, 19(4):659–671.

Jacinto, L. R., Reis, J. S., Dias, N. S., Cerqueira, J. J., Correia, J. H., and Sousa, N. (2013). Stress affects theta activity in limbic networks and impairs novelty-induced exploration and familiarization. Frontiers in Behavioral Neuroscience, 7:127.

Jenkinson, M., Bannister, P., Brady, M., and Smith, S. (2002). Improved optimization for the robust and accurate linear registration and motion correction of brain images. Neuroimage, 17(2):825–841.

Jenkinson, M. and Smith, S. (2001). A global optimisation method for robust affine registration of brain images. Medical image analysis, 5(2):143–156.

Joëls, M. and Baram, T. Z. (2009). The neuro-symphony of stress. Nature reviews neuroscience, 10(6):459–466.

Joëls, M., Pu, Z., Wiegert, O., Oitzl, M. S., and Krugers, H. J. (2006). Learning under stress: how does it work? Trends in cognitive sciences, 10(4):152–158.

Kamp, S.-M., Endemann, R., Domes, G., and Mecklinger, A. (2019). Effects of acute psychosocial stress on the neural correlates of episodic encoding: Item versus associative memory. Neurobiology of Learning and Memory, 157:128–138.

Karst, H., Berger, S., Turiault, M., Tronche, F., Schütz, G., and Joëls, M. (2005). Mineralocorticoid receptors are indispensable for nongenomic modulation of hippocampal glutamate transmission by corticosterone. Proceedings of the National Academy of Sciences, 102(52):19204–19207.

Karst, H. and Joëls, M. (2005). Corticosterone slowly enhances miniature excitatory postsynaptic current amplitude in mice ca1 hippocampal cells. Journal of Neurophysiology, 94(5):3479–3486.

Kim, J. J. and Diamond, D. M. (2002). The stressed hippocampus, synaptic plasticity and lost memories. Nature Reviews Neuroscience, 3(6):453–462.

Kim, J. J., Lee, H. J., Han, J.-S., and Packard, M. G. (2001). Amygdala is critical for stress-induced modulation of hippocampal long-term potentiation and learning. Journal of Neuroscience, 21(14):5222–5228.

Krugers, H. J., Alfarez, D. N., Karst, H., Parashkouhi, K., van Gemert, N., and Joëls, M. (2005). Corticosterone shifts different forms of synaptic potentiation in opposite directions. Hippocampus, 15(6):697–703.

Kuhlmann, S. and Wolf, O. T. (2006). Arousal and cortisol interact in modulating memory consolidation in healthy young men. Behavioral neuroscience, 120(1):217.

LaRocque, K. F., Smith, M. E., Carr, V. A., Witthoft, N., Grill-Spector, K., and Wagner, A. D. (2013). Global similarity and pattern separation in the human medial temporal lobe predict subsequent memory. Journal of Neuroscience, 33(13):5466–5474.

Lenth, R. V. (2022). emmeans: Estimated Marginal Means, aka Least-Squares Means. R package version 1.7.5.

Lovallo, W. R., Robinson, J. L., Glahn, D. C., and Fox, P. T. (2010). Acute effects of hydrocortisone on the human brain: an fmri study. Psychoneuroendocrinology, 35(1):15–20.

Lupien, S. J., Maheu, F., Tu, M., Fiocco, A., and Schramek, T. E. (2007). The effects of stress and stress hormones on human cognition: Implications for the field of brain and cognition. Brain and Cognition, 65(3):209–237.

Madan, C. R., Scott, S. M., and Kensinger, E. A. (2019). Positive emotion enhances association-memory. Emotion, 19(4):733.

Mäkelä, K. (2004). Studies of the reliability and validity of the addiction severity index. Addiction, 99(4):398–410.

McClelland, J. L., McNaughton, B. L., and O’Reilly, R. C. (1995). Why there are complementary learning systems in the hippocampus and neocortex: insights from the successes and failures of connectionist models of learning and memory. Psychological review, 102(3):419.

McEwen, B. S. and Sapolsky, R. M. (1995). Stress and cognitive function. Current opinion in neurobiology, 5(2):205–216.

McGaugh, J. L. et al. (2015). Consolidating memories. Annu Rev Psychol, 66(1):1–24.

Meier, J. K., Weymar, M., and Schwabe, L. (2020). Stress alters the neural context for building new memories. Journal of Cognitive Neuroscience, 32(12):2226–2240.

Molitor, R. J., Sherrill, K. R., Morton, N. W., Miller, A. A., and Preston, A. R. (2021). Memory reactivation during learning simultaneously promotes dentate gyrus/ca2, 3 pattern differentiation and ca1 memory integration. Journal of Neuroscience, 41(4):726–738.

Nolan, M., Roman, E., Nasa, A., Levins, K. J., O’Hanlon, E., O’Keane, V., and Willian Roddy, D. (2020). Hippocampal and amygdalar volume changes in major depressive disorder: a targeted review and focus on stress. Chronic Stress, 4:2470547020944553.

Norman-Haignere, S. V., McCarthy, G., Chun, M. M., and Turk-Browne, N. B. (2012). Category-selective background connectivity in ventral visual cortex. Cerebral Cortex, 22(2):391–402.

Okuda, S., Roozendaal, B., and McGaugh, J. L. (2004). Glucocorticoid effects on object recognition memory require training-associated emotional arousal. Proceedings of the National Academy of Sciences, 101(3):853–858.

Patenaude, B., Smith, S. M., Kennedy, D. N., and Jenkinson, M. (2011). A bayesian model of shape and appearance for subcortical brain segmentation. Neuroimage, 56(3):907–922.

Pavlides, C., Watanabe, Y., Magarin, A., McEwen, B., et al. (1995). Opposing roles of type i and type ii adrenal steroid receptors in hippocampal long-term potentiation. Neuroscience, 68(2):387–394.

Payne, J., Jackson, E., Ryan, L., Hoscheidt, S., Jacobs, J., and Nadel, L. (2006). The impact of stress on neutral and emotional aspects of episodic memory. Memory, 14(1):1–16.

Payne, J. D., Jackson, E. D., Hoscheidt, S., Ryan, L., Jacobs, W. J., and Nadel, L. (2007). Stress administered prior to encoding impairs neutral but enhances emotional long-term episodic memories. Learning & Memory, 14(12):861–868.

Pedraza, L. K., Sierra, R. O., Boos, F. Z., Haubrich, J., Quillfeldt, J. A., and de Oliveira Alvares, L. (2016). The dynamic nature of systems consolidation: Stress during learning as a switch guiding the rate of the hippocampal dependency and memory quality. Hippocampus, 26(3):362–371.

Pelli, D. G. (1997). The videotoolbox software for visual psychophysics: transforming numbers into movies. Spatial Vision.

Pinheiro, J., Bates, D., and R Core Team (2022). nlme: Linear and Nonlinear Mixed Effects Models. R package version 3.1-161.

Pruessner, J. C., Dedovic, K., Khalili-Mahani, N., Engert, V., Pruessner, M., Buss, C., Renwick, R., Dagher, A., Meaney, M. J., and Lupien, S. (2008). Deactivation of the limbic system during acute psychosocial stress: evidence from positron emission tomography and functional magnetic resonance imaging studies. Biological psychiatry, 63(2):234–240.

Pruessner, J. C., Köhler, S., Crane, J., Pruessner, M., Lord, C., Byrne, A., Kabani, N., Collins, D. L., and Evans, A. C. (2002). Volumetry of temporopolar, perirhinal, entorhinal and parahippocampal cortex from high-resolution mr images: considering the variability of the collateral sulcus. Cerebral Cortex, 12(12):1342–1353.

Qin, S., Hermans, E. J., Van Marle, H. J., and Fernández, G. (2012). Understanding low reliability of memories for neutral information encoded under stress: alterations in memory-related activation in the hippocampus and midbrain. Journal of Neuroscience, 32(12):4032–4041.

Rey, M., Carlier, E., Talmi, M., and Soumireu-Mourat, B. (1994). Corticosterone effects on long-term potentiation in mouse hippocampal slices. Neuroendocrinology, 60(1):36–41.

Roozendaal, B., McEwen, B. S., and Chattarji, S. (2009). Stress, memory and the amygdala. Nature Reviews Neuroscience, 10(6):423–433.

Roozendaal, B. and McGaugh, J. L. (1997). Basolateral amygdala lesions block the memory-enhancing effect of glucocorticoid administration in the dorsal hippocampus of rats. European Journal of Neuroscience, 9(1):76–83.

Roozendaal, B., Okuda, S., Van der Zee, E. A., and McGaugh, J. L. (2006). Glucocorticoid enhancement of memory requires arousal-induced noradrenergic activation in the basolateral amygdala. Proceedings of the National Academy of Sciences, 103(17):6741–6746.

Sandi, C. and Pinelo-Nava, M. T. (2007). Stress and memory: behavioral effects and neurobiological mechanisms. Neural plasticity, 2007.

Schapiro, A. C., Turk-Browne, N. B., Botvinick, M. M., and Norman, K. A. (2017). Complementary learning systems within the hippocampus: a neural network modelling approach to reconciling episodic memory with statistical learning. Philosophical Transactions of the Royal Society B: Biological Sciences, 372(1711):20160049.

Schwabe, L., Bohringer, A., Chatterjee, M., and Schachinger, H. (2008). Effects of pre-learning stress on memory for neutral, positive and negative words: Different roles of cortisol and autonomic arousal. Neurobiology of learning and memory, 90(1):44–53.

Schwabe, L., Hermans, E. J., Joëls, M., and Roozendaal, B. (2022). Mechanisms of memory under stress. Neuron.

Seckl, J. R., Dickson, K. L., Yates, C., and Fink, G. (1991). Distribution of glucocorticoid and mineralocorticoid receptor messenger rna expression in human postmortem hippocampus. Brain Research, 561(2):332–337.

Segal, M., Richter-Levin, G., and Maggio, N. (2010). Stress-induced dynamic routing of hippocampal connectivity: A hy-pothesis.

Segal, S., Simon, R., McFarlin, S., Alkire, M., Desai, A., and Cahill, L. (2014). Glucocorticoids interact with noradrenergic activation at encoding to enhance long-term memory for emotional material in women. Neuroscience, 277:267–272.

Sharvit, A., Segal, M., Kehat, O., Stork, O., and Richter-Levin, G. (2015). Differential modulation of synaptic plasticity and local circuit activity in the dentate gyrus and ca1 regions of the rat hippocampus by corticosterone. Stress, 18(3):319–327.

Shields, G. S., Hunter, C. L., and Yonelinas, A. P. (2022). Stress and memory encoding: What are the roles of the stress-encoding delay and stress relevance? Learning & Memory, 29(2):48–54.

Shields, G. S., Sazma, M. A., McCullough, A. M., and Yonelinas, A. P. (2017). The effects of acute stress on episodic memory: A meta-analysis and integrative review. Psychological bulletin, 143(6):636.

Shors, T. J. (2006). Stressful experience and learning across the lifespan. Annual review of psychology, 57:55.

Sinha, R., Fogelman, N., Wemm, S., Angarita, G., Seo, D., and Hermes, G. (2022). Alcohol withdrawal symptoms predict cor-ticostriatal dysfunction that is reversed by prazosin treatment in alcohol use disorder. Addiction Biology, 27(2):e13116.

Sinha, R., Lacadie, C. M., Constable, R. T., and Seo, D. (2016). Dynamic neural activity during stress signals resilient coping. Proceedings of the National Academy of Sciences, 113(31):8837–8842.

Smeets, T., Giesbrecht, T., Jelicic, M., and Merckelbach, H. (2007). Context-dependent enhancement of declarative mem-ory performance following acute psychosocial stress. Biological psychology, 76(1-2):116–123.

Smith, S. M. (2002). Fast robust automated brain extraction. Human brain mapping, 17(3):143–155.

Stepan, J., Dine, J., Fenzl, T., Polta, S. A., von Wolff, G., Wotjak, C. T., and Eder, M. (2012). Entorhinal theta-frequency input to the dentate gyrus trisynaptically evokes hippocampal ca1 ltp. Frontiers in neural circuits, 6:64.

Symonds, C. S., McKie, S., Elliott, R., Deakin, J. F. W., and Anderson, I. M. (2012). Detection of the acute effects of hydro-cortisone in the hippocampus using pharmacological fmri. European Neuropsychopharmacology, 22(12):867–874.

Tambini, A., Rimmele, U., Phelps, E. A., and Davachi, L. (2017). Emotional brain states carry over and enhance future memory formation. Nature neuroscience, 20(2):271–278.

Tompary, A. and Davachi, L. (2017). Consolidation promotes the emergence of representational overlap in the hippocam-pus and medial prefrontal cortex. Neuron, 96(1):228–241.

Ulrich-Lai, Y. M. and Herman, J. P. (2009). Neural regulation of endocrine and autonomic stress responses. Nature reviews neuroscience, 10(6):397–409.

Vaisvaser, S., Lin, T., Admon, R., Podlipsky, I., Greenman, Y., Stern, N., Fruchter, E., Wald, I., Pine, D. S., Tarrasch, R., et al. (2013). Neural traces of stress: cortisol related sustained enhancement of amygdala-hippocampal functional connectivity. Frontiers in human neuroscience, 7:313.

van Ast, V. A., Cornelisse, S., Meeter, M., and Kindt, M. (2014). Cortisol mediates the effects of stress on the contextual dependency of memories. Psychoneuroendocrinology, 41:97–110.

Van Der Linden, L., Mathôt, S., and Vitu, F. (2015). The role of object affordances and center of gravity in eye movements toward isolated daily-life objects. Journal of Vision, 15(5):8–8.

van Stegeren, A. H. (2009). Imaging stress effects on memory: a review of neuroimaging studies. The Canadian Journal of Psychiatry, 54(1):16–27.

Vandael, D., Wierda, K., Vints, K., Baatsen, P., De Groef, L., Moons, L., Rybakin, V., and Gounko, N. V. (2021). Corticotropin-releasing factor induces functional and structural synaptic remodelling in acute stress. Translational Psychiatry, 11(1):1–12.

Visser, R. M., Scholte, H. S., Beemsterboer, T., and Kindt, M. (2013). Neural pattern similarity predicts long-term fear memory. Nature neuroscience, 16(4):388–390.

Wammes, J., Norman, K. A., and Turk-Browne, N. (2022). Increasing stimulus similarity drives nonmonotonic representa-tional change in hippocampus. ELife, 11.

Wang, Q., Van Heerikhuize, J., Aronica, E., Kawata, M., Seress, L., Joels, M., Swaab, D. F., and Lucassen, P. J. (2013). Glucocor-ticoid receptor protein expression in human hippocampus; stability with age. Neurobiology of Aging, 34(6):1662–1673.

Wang, Z., Neylan, T. C., Mueller, S. G., Lenoci, M., Truran, D., Marmar, C. R., Weiner, M. W., and Schuff, N. (2010). Magnetic resonance imaging of hippocampal subfields in posttraumatic stress disorder. Archives of general psychiatry, 67(3):296–303.

Wanjia, G., Favila, S. E., Kim, G., Molitor, R. J., and Kuhl, B. A. (2021). Abrupt hippocampal remapping signals resolution of memory interference. Nature communications, 12(1):1–11.

Watson, D., Clark, L. A., and Tellegen, A. (1988). Development and validation of brief measures of positive and negative affect: the panas scales. Journal of personality and social psychology, 54(6):1063.

Weis, C. N., Webb, E. K., Huggins, A. A., Kallenbach, M., Miskovich, T. A., Fitzgerald, J. M., Bennett, K. P., Krukowski, J. L., deRoon Cassini, T. A., and Larson, C. L. (2021). Stability of hippocampal subfield volumes after trauma and relationship to development of ptsd symptoms. NeuroImage, 236:118076.

Wiegert, O., Joëls, M., and Krugers, H. (2006). Timing is essential for rapid effects of corticosterone on synaptic potentiation in the mouse hippocampus. Learning & Memory, 13(2):110–113.

Woolrich, M. W., Ripley, B. D., Brady, M., and Smith, S. M. (2001). Temporal autocorrelation in univariate linear modeling of fmri data. Neuroimage, 14(6):1370–1386.

Xiao, J., Hays, J., Ehinger, K. A., Oliva, A., and Torralba, A. (2010). Sun database: Large-scale scene recognition from abbey to zoo. In 2010 IEEE computer society conference on computer vision and pattern recognition, pages 3485–3492. IEEE.

Yushkevich, P. A., Pluta, J. B., Wang, H., Xie, L., Ding, S.-L., Gertje, E. C., Mancuso, L., Kliot, D., Das, S. R., and Wolk, D. A. (2015). Automated volumetry and regional thickness analysis of hippocampal subfields and medial temporal cortical structures in mild cognitive impairment. Human brain mapping, 36(1):258–287.

Zoladz, P. R., Clark, B., Warnecke, A., Smith, L., Tabar, J., and Talbot, J. N. (2011). Pre-learning stress differentially affects long-term memory for emotional words, depending on temporal proximity to the learning experience. Physiology & behavior, 103(5):467–476.

