## Supplemental Figures 1-4 for "Hippocampal mechanisms support cortisol-induced memory enhancements"

**Supplemental Information for: Hippocampal mechanisms support cortisol-induced memory enhancements**

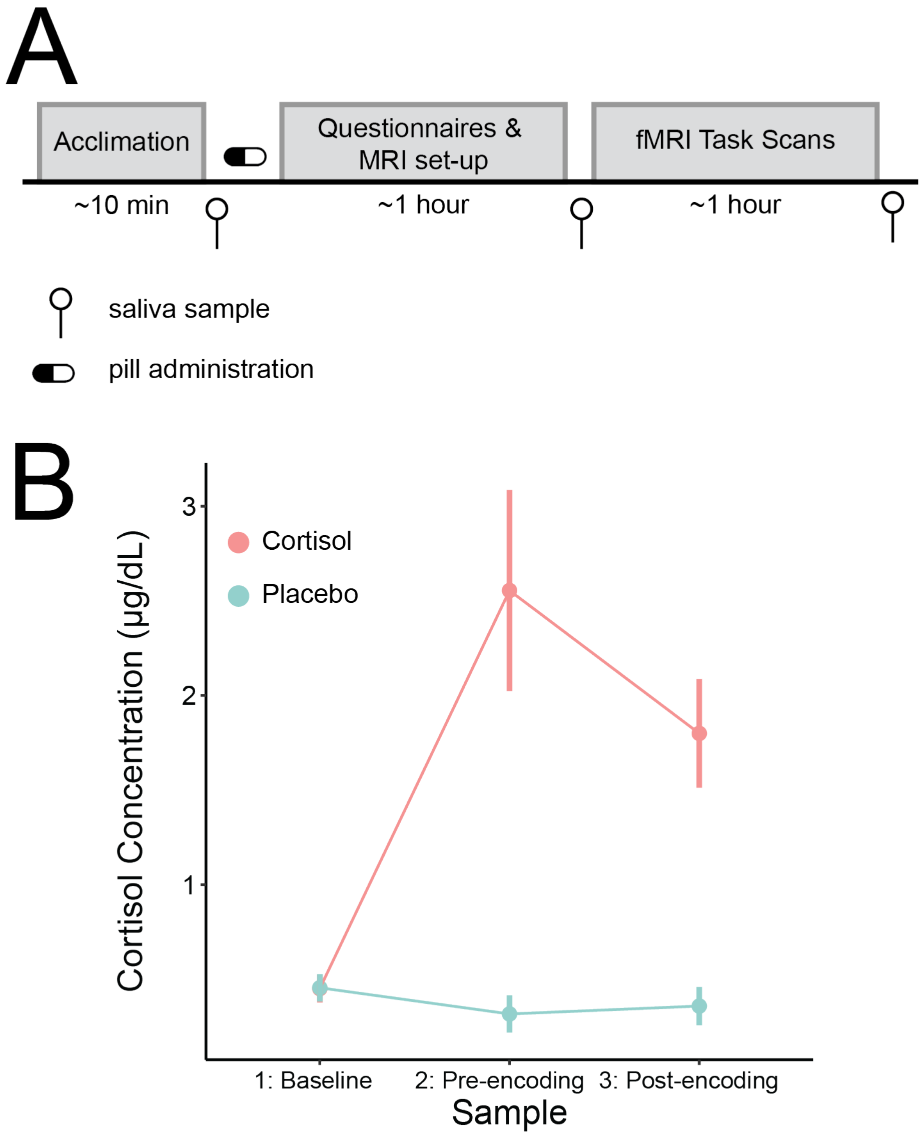


**Fig. S1.** Salivary cortisol measurements. A) Participants provided three saliva samples throughout the course of the encoding session: prior to pill administration (after a 10-minute acclimation period), prior to encoding (approximately 1 hour post-pill), and after encoding (approximately 2 hours post-pill). B) Hydrocortisone led to elevated salivary cortisol concentrations, as reflected by a main effect of pill [$F(1, 118) = 35.12, p < 0.001$], a main effect of timepoint [$F(2, 118) = 7.27, p = 0.001$], and a pill x timepoint interaction [$F(2, 118) = 9.38, p < 0.001$]. Specifically, salivary cortisol was elevated at the two post-pill administration timepoints, indicating that cortisol was elevated during memory encoding.

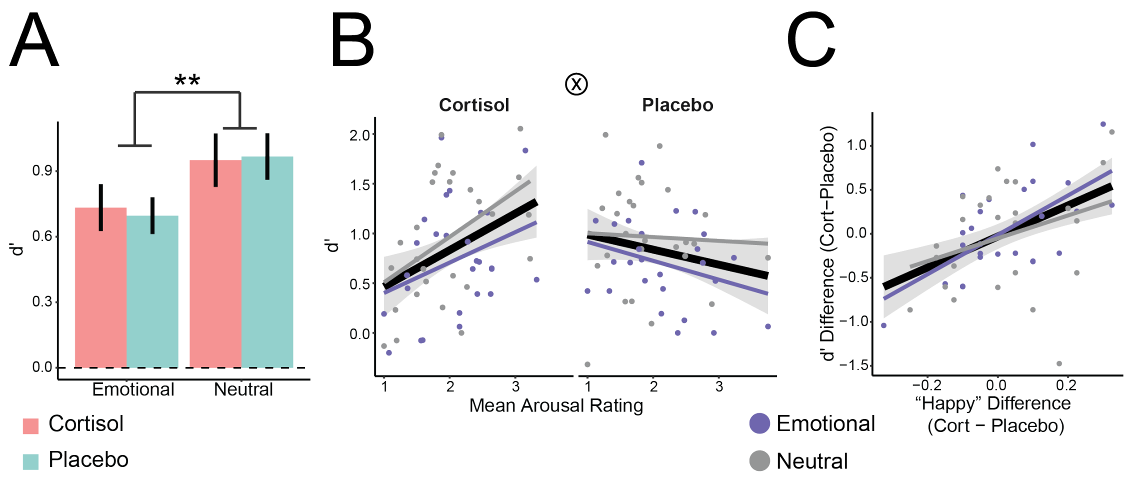


**Fig. S2.** Object Recognition Results. A) Participants exhibited above chance memory for individual objects, as indicated by d' >0 (dashed lined; Mean d' = 0.84; SD = 0.43). There was a main effect of trial type on d' [$F(1, 75) = 10.60, p = 0.0017$], with better memory for neutral objects. B) There was also an interaction between arousal and pill on memory [$F(1, 71) = 10.36, p = 0.0019$], such that participants with higher subjective arousal ratings had better object memory under hydrocortisone, but worse memory under placebo. C) The change in participants' subjective "happy" ratings predicted the difference in memory performance from hydrocortisone to placebo, [$F(1, 23) = 12.08, p = 0.0020$], with more "happy" ratings under hydrocortisone corresponding to better memory. A: Error bars indicate standard error of the mean across participants; **p <0.01. B-C: Error shading indicates 95% confidence interval around the line of best fit, collapsed across trial types (black line). Individual colored lines indicate the line of best fit per trial type.

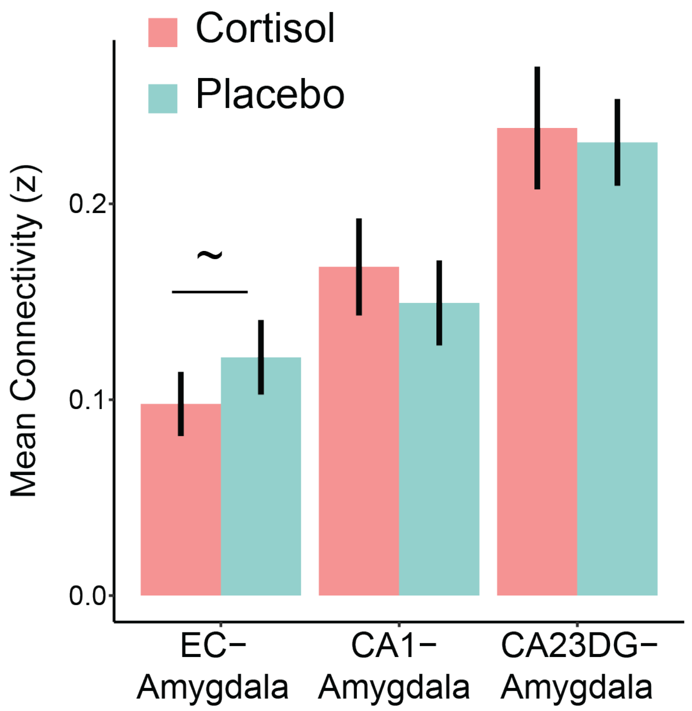


**Fig. S3.** Hippocampal-amygdala background connectivity results. Error bars indicate standard error of the mean across participants; ~p<0.09.

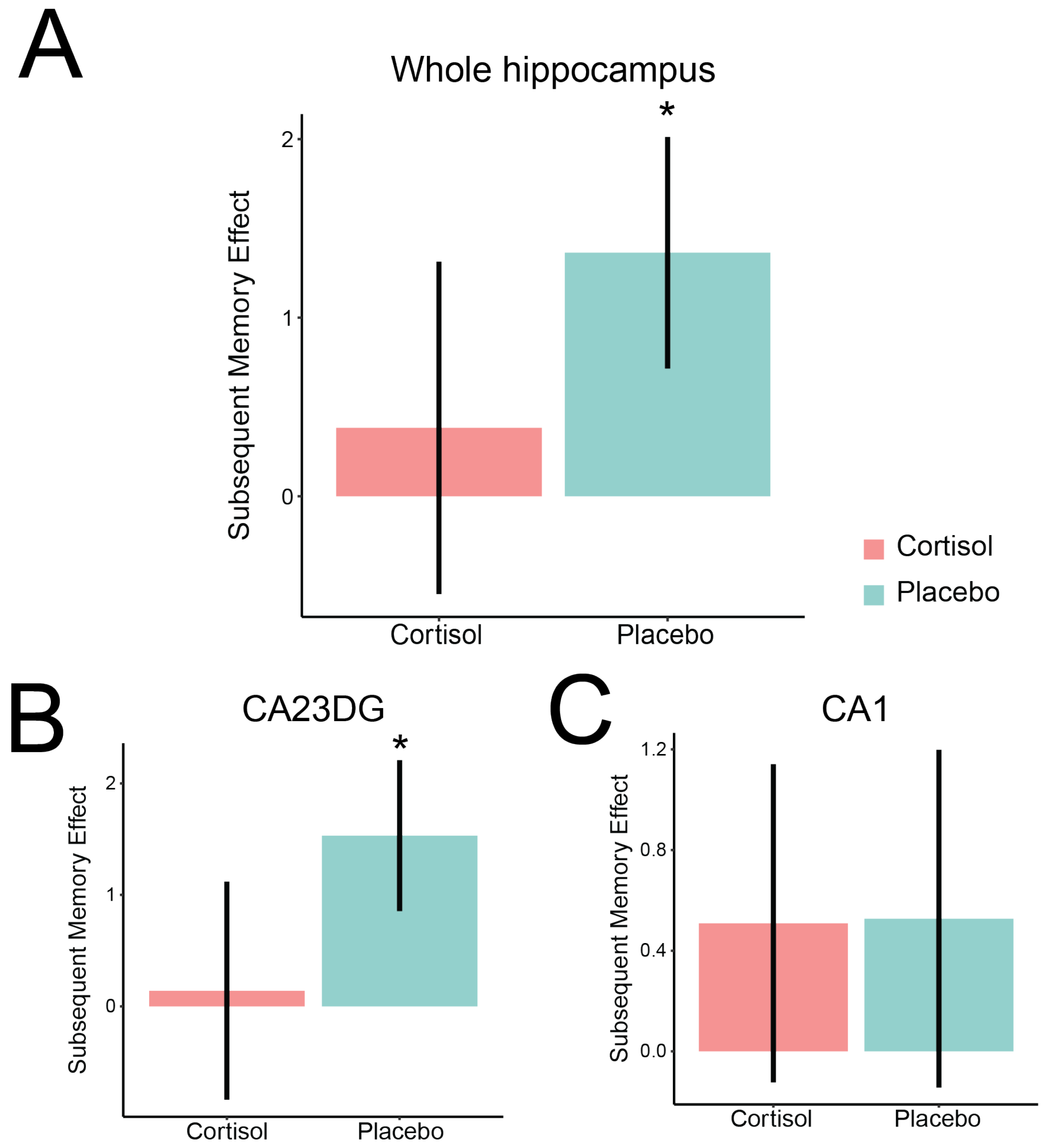


**Fig. S4.** Hippocampal subsequent memory effects. A) Univariate subsequent memory effect (difference in encoding activation for subsequently remembered, versus forgotten trials), averaged across all voxels in the hippocampus. B) We observed a similar pattern within CA23DG: although there were no main effects of pill or trial type, nor interactions ($ps > 0.20$), there was a reliable subsequent memory effect under placebo [Mean = 1.53; SD = 3.38; $t(24) = 2.26, p = 0.033$] but not hydrocortisone [Mean = 0.14; SD = 4.99; $t(25) = 0.14, p = 0.89$]. C) In CA1, we again did not observe main effects of pill or trial type, nor interactions ($ps > 0.30$), but (in contrast to CA23DG) we did not observe any reliable subsequent memory effects [placebo: Mean = 0.53; SD = 3.36; $t(24) = 0.79, p = 0.44$; hydrocortisone: Mean = 0.51; SD = 3.22; $t(25) = 0.80, p = 0.43$]. Error bars indicate standard error of the mean across participants; *p <0.05.
